# A Quasi Birth-and-Death Model For Tumor Recurrence

**DOI:** 10.1101/551770

**Authors:** Leonardo M. Santana, Shridar Ganesan, Gyan Bhanot

**Affiliations:** Department of Physics & Astronomy, Rutgers University, Piscataway, New Jersey 08854, USA; Rutgers Cancer Institute of New Jersey, New Brunswick, New Jersey 08903, USA; Department of Medicine, Rutgers Robert Wood Johnson Medical School, Rutgers University, New Brunswick, New Jersey 08903, USA; Department of Molecular Biology and Biochemistry, Rutgers University, Piscataway, New Jersey 08854, USA

**Keywords:** Survival analysis, Tumor growth dynamics, Dormant tumor, Master equation, First-passage time

## Abstract

A major cause of chemoresistance and recurrence in tumors is the presence of dormant tumor foci that survive chemotherapy and can eventually transition to active growth to regenerate the cancer. In this paper, we propose a Quasi Birth-and-Death (QBD) model for the dynamics of tumor growth and recurrence/remission of the cancer. Starting from a discrete-state master equation that describes the time-dependent transition probabilities between states with different numbers of dormant and active tumor foci, we develop a framework based on a continuum-limit approach to determine the time-dependent probability that an undetectable residual tumor will become large enough to be detectable. We derive an exact formula for the probability of recurrence at large times and show that it displays a phase transition as a function of the ratio of the death rate *µ*_*A*_ of an active tumor focus to its doubling rate *λ*. We also derive forward and backward Kolmogorov equations for the transition probability density in the continuum limit and, using a first-passage time formalism, we obtain a drift-diffusion equation for the mean recurrence time and solve it analytically to leading order for a large detectable tumor size *N*. We show that simulations of the discrete-state model agree with the analytical results, except for *O*(1*/N*) corrections. Finally, we describe a scheme to fit the model to recurrence-free survival (Kaplan-Meier) curves from clinical cancer data, using ovarian cancer data as an example. Our model has potential applications in predicting how changing chemotherapy schedules may affect disease recurrence rates, especially in cancer types for which no targeted therapy is available.

## 1. Introduction

The advent of chemotherapy was an important milestone in the history of cancer treatment and research. For the treatment of early stage cancer, it remains the only option after surgery and radiation for cancers where no long-term targeted adjuvant (post-surgery) therapy is available, such as serous ovarian cancer and triple negative (ER-/HER2-/PR-) breast cancer [1]. By targeting rapidly cycling cells, chemotherapy systemically attacks growing tumors. Side effects on other tissues, especially on cells with a high turnover rate, such as skin and the intestinal epithelium, can be moderate to severe, depending on the duration and intensity of treatment.

Several mathematical models have been proposed to predict optimal regimens of adjuvant chemotherapy to specify duration, dosage levels or dosing protocols, with the goal of reducing recurrence hazard rates. These models may be classified by their mathematical approach, which can be either stochastic or deterministic and linear or nonlinear, and to the nature of the biological assumptions underlying them (see [2] for a review). Examples of such models include optimal-control-theory models [3], game-theoretical models [4], as well as models of drug resistance and/or chemotherapy scheduling, which can be either stochastic [5–8] or deterministic [9–11].

In current clinical practice, chemotherapy is usually given at “maximum tolerated dose” for the “minimum possible duration”, which is usually 3-6 months, in the belief that this will have the most benefit to the patient in the least possible time [13]. This is based on modeling tumor cells as continuously dividing at some fixed deterministic rate [14]. Norton and Simon [15] proposed that tumor growth follows a type of sigmoid curve known in the literature as the Gompertzian function, and also proposed the tumor-regression hypothesis, that cell kill is proportional to the growth rate of the untreated tumor [12, 16]. Since the Gompertzian growth rate decreases as the tumor grows, they concluded that it becomes increasingly more difficult to kill the tumor as its size increases. This model provides the justification for the “maximum tolerated dose” and maximizing dose density, the goal being to efficiently kill the tumor when it is small and growing rapidly. However, the effectiveness of this treatment paradigm is unclear and not entirely consistent with some clinical and experimental data [17, 18] (see also the review paper [16] for a detailed literature review).

Furthermore, it is known that the *stage* of a cancer, which is related to the tumor size and degree of lymph node involvement, is an excellent predictor of prognosis, independently of cancer type or therapy. The larger the tumor, the more difficult it is to effect a cure. Likewise, tumor *grade*, which is a measure of tumor aggressiveness, is also a good predictor of outcome. However, it has been suggested that the fractional impact of treatment on improved survival is higher for patients with late-stage or high-grade tumors than for patients with early-stage or low-grade tumors (see, for example, the studies [20] and [21]). This observation argues against the high-dose/short-term treatment paradigm by suggesting that unlike Norton and Simon proposed in their deterministic Gompertzian growth model, tumor cells may not all be in continuous growth, and that actively growing tumors tend to be more responsive to cytotoxic drugs than those that are mostly in a dormant or resting state.

In [19], a stochastic alternative to Norton and Simon’s deterministic Gompertzian model is proposed for breast cancer, where tumors are not in continuous growth, but can be either in a dormant state or in an active-growth state. Indeed, it has been suggested that a major cause of resistance to chemotherapy is the presence of dormant tumor foci with a cycle time that exceeds the duration of chemotherapy [18, 19, 22]. This is consistent with the observation that often, the effect of chemotherapy on recurrence rates does not last for a long time after chemotherapy ends. Several clinical trials [23–25] have shown that improved recurrence rates for patients receiving chemotherapy revert to rates for the control group (who received no chemotherapy) in a relatively short time after termination of treatment, suggesting the presence of residual dormant tumor foci that survive treatment and regenerate the cancer. These data also suggest that short-duration chemotherapy only targets the tumor foci that cycle during chemotherapy. Tumor foci that cycle after chemotherapy ends are not affected and can cause recurrence. This suggests that chemotherapy may be more effective if its duration is optimized to the time it takes dormant foci to transition to active growth. Direct evidence for this hypothesis is available in analysis of lymphoma data [26, 27], where maximizing chemotherapy dosage did not have a prolonged effect on outcome, whereas extending the duration of chemotherapy, while maintaining a minimum effective dose was more beneficial. Long-term hormonal therapy with drugs that target the ER pathway (e.g. tamoxifen) in ER+/HER2-breast cancers, which are low grade (have low transition rates of tumor foci from dormancy to active growth) provides yet another example that longer-term treatment is preferable to short-term treatment [28]. These studies provide evidence that optimally adjusting the duration of chemotherapy may improve chemotherapy effectiveness, while maintaining the same total amount of drug administered over the course of the chemotherapy regimen.

These observations suggest the following two hypotheses: 1) For a given cancer type, there is a characteristic time for dormant tumor foci to transition to active growth; 2) Dormant tumor foci are often resistant to chemotherapeutic agents, while active tumor foci are not. Based on these hypotheses, we develop a mathematical model and framework with potential application to study the impact of variation in dosage and duration of chemotherapy on recurrence rates. Our model may serve as a guide for the design of experiments and clinical trials that may eventually lead to optimized chemotherapy regimens.

The remainder of the paper is organized as follows. In Section 2, we give an overview of the model by defining its state space, parameters and transition rules. In Section 3, the stochastic dynamics of the model is formulated in terms of a continuous-time master equation in the discrete state space that represents the numbers of dormant and active tumor foci. In Section 4, it is shown that an expansion of the master equation for a large detectable-tumor size *N* leads to a simplified approach by mapping the original discrete-state model to a stochastic process in a continuous two-dimensional state space. In Section 5, we find the large-time probability of recurrence in closed analytic form and calculate the mean recurrence time (MRT) analytically to leading order in *N*. In Section 5, we compare these analytical results to simulations and describe a scheme to fit the model to recurrence-free survival (Kaplan-Meier) curves from clinical cancer data, using ovarian cancer data as an example. Finally, in Section (6) we present our concluding remarks.

## 2. Overview of the discrete-state model

The precise discrete model for tumor recurrence that will be described in this section was inspired by previous work on the effect of quiescence (i.e., the presence of dormant tumor foci) on treatment success, such as the work of Komarova and Wodarz [8], which inspired the stochastic model described below, or to deterministic versions of their model, proposed in [9], [10], or [11]. However, in contrast to these earlier studies, the focus of our paper is on finding a relationship between the model parameters and the time to recurrence for a given treatment regimen. This is achieved using special boundary conditions that will be described in Sections 3 and 4, which to our knowledge, is a novel contribution to tumor modeling. For simplicty and as a first exercise, we will explore only treatment regimens controlled by a single parameter *µ*_*A*_, the death rate of actively-dividing tumor foci. Also, unlike previous work mentioned above, we will only model transitory chemoresistance resulting from dormancy, without discussing the important but more difficult stochastic issue of resistance from acquired mutations.

After surgery and radiation therapy, we assume that cancer patients retain a number of residual, undetectable tumor foci that may eventually grow and create a detectable tumor (recurrence). The foci can transition from a dormant (non-dividing, chemo-resistant) state to an active (dividing, chemo-sensitive) state and vice versa with rates *η* and *ξ* respectively. Chemotherapy affects only active foci, which can either double or die, with rates *λ* and *µ*_*A*_ respectively. The dynamics of this process is modeled as a Quasi Birth-and-Death (QBD) process (for an introduction to the topic, see [29, 30]) that describes the stochastic time evolution of the number of active and dormant tumor foci in a patient, resulting in either tumor recurrence (when the number of foci is large enough to be detected) or remission (when there are no foci left).

Denoting by *|m, n*〉 the state with *m* dormant (𝒟) tumor foci and *n* active (𝒜) foci, the goal is to predict the time evolution of the joint probability distribution *p*_*mn*_(*t*), given the initial condition *p*_*mn*_(0). Without loss of generality, we can choose *p*_*mn*_(0) = *δ*_*m*0,*n*0_, where *m*_0_ and *n*_0_ are respectively initial counts of dormant and active foci. This is so because the latter initial condition defines a Green’s function, from which the solution for any arbitrary initial condition can be constructed as a convolution of transition probabilities, because of linearity.

Our QBD model for tumor recurrence is a Markov process on the two-dimensional lattice of all possible Fock states *|m, n〉* with the following transition rules:

- At any given time, an active (𝒜) tumor focus may either double or die at rates *λ* and *µ*_*A*_, respectively. During chemotherapy (between time *t* = 0 and time *t* = *t*_*chemo*_ > 0), the death rate is *µ*_*A*_ = *µ*_*chemo*_ and after treatment (beyond time *t* = *t*_*chemo*_), it decreases to a lower baseline rate *µ*_*A*_ = *µ*_0_.
- By definition, a dormant (𝒟) tumor focus cannot double, but it could in principle die at some rate *µ*_*D*_. However, we will eventually set *µ*_*D*_ = 0. This is because dormant foci can repair chemotherapy damage as we argued above, i.e., they are chemoresistant.
- An active (𝒜) tumor focus may transition to dormancy (𝒟) at rate *ξ* and a dormant tumor focus may become active at rate *η*.
- Let 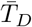 and 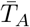 be the respective times that a tumor focus spends, on average, in the dormant and active states before it doubles (averaged over many doublings). It then follows that the 𝒟 → 𝒜 and 𝒜 → 𝒟 hopping rates are given by 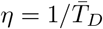 and 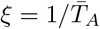, respectively, so the doubling rate is given by 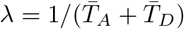. Hence, the rates *η*, *ξ* and *λ* are related by the equation

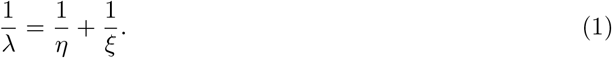 This constraint reduces the number of parameters in the model by one and allows us to parametrize the rates *η*, *ξ* as

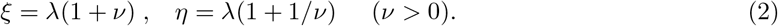
- All the tumor foci within each population (either 𝒟 or 𝒜) are equally likely to undergo a transition, so each transition probability on the two-dimensional lattice of states is proportional to either the population of dormant foci (*m*) or the population of active foci (*n*). The transition probabilities of any transitions beyond nearest neighbors are assumed to be second order in infinitesimal time, so the model is “skip free”. The transition probabilities to neighboring states are then given by

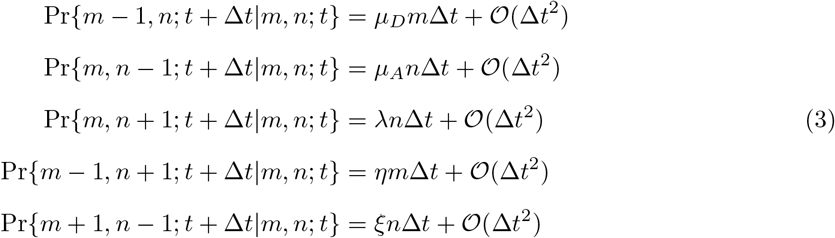

and for transitions to states beyond nearest-neighbors, i.e., for |*m* − *m*′| > 1 or |*n* − *n*′| > 1, we assume

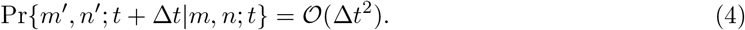 A diagram of the state space showing the transitions above is given in Fig. 1.
- If the total number of tumor foci *m* + *n* reaches a sufficiently large number *N*, the tumor becomes detectable and no further transitions are allowed, i.e., the disease recurrence is defined by means of absorbing boundary conditions at the recurrence boundary *m* + *n* = *N*. The absorbing boundary condition at the extinction state (0, 0) is automatically satisfied, since the transition rates are proportional to either *m* or *n*.

**Figure 1:**
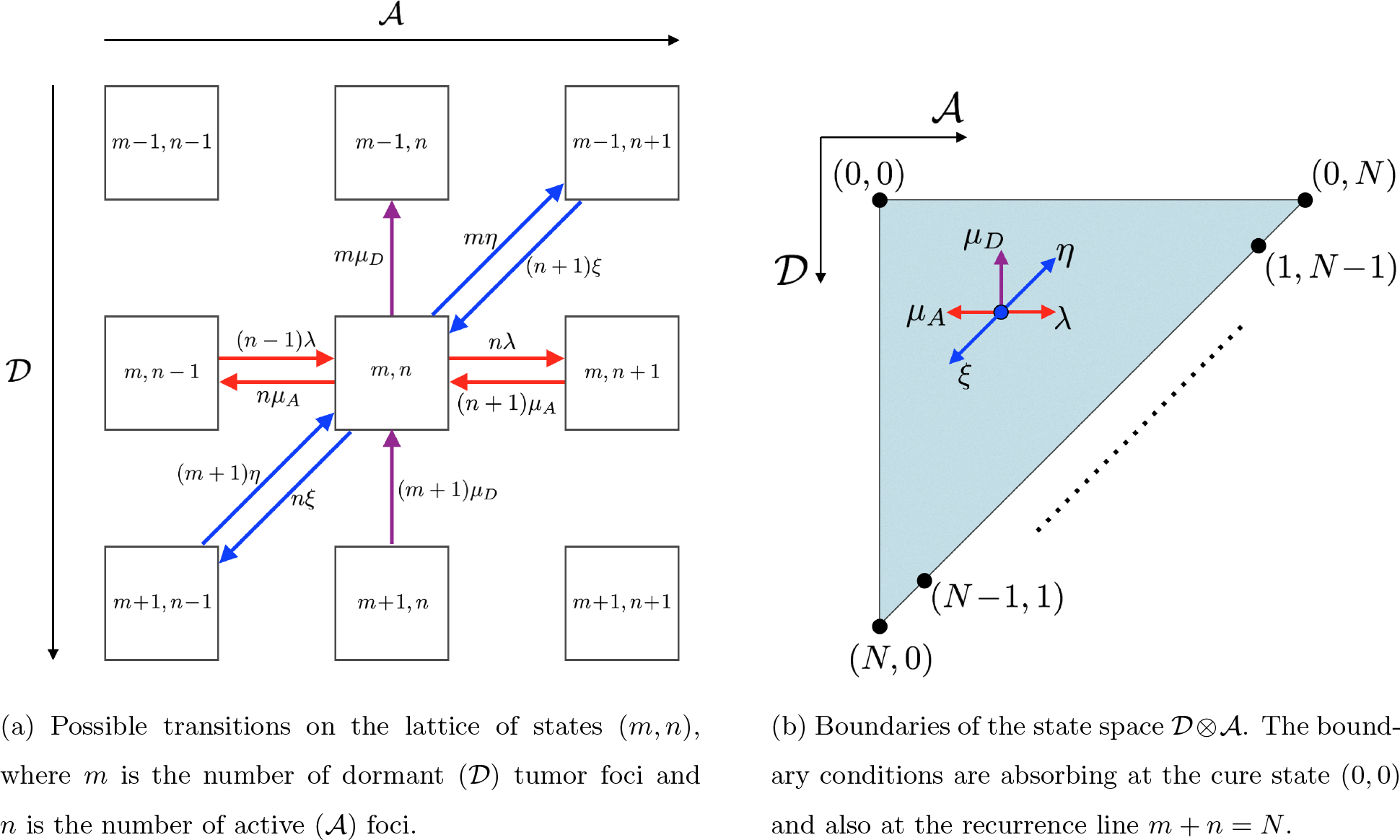
Structure of the Fock-like state space of the QBD model for tumor recurrence.

## 3. Master equation for the state probabilities

From the transition rules described in Section 2, it follows that the time evolution of the state probabilities *p*_*mn*_(*t*) is given by the master equation

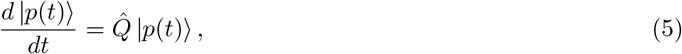

where |*p*(*t*)〉 is the probability vector, whose components are the state probabilities *p*_*mn*_(*t*), i.e., |*p*(*t*)〉 = 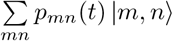. The infinitesimal transition operator 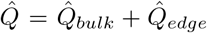 consists of a “bulk” part 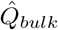 and an edge correction 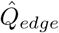 due to the absorbing boundary conditions at the recurrence line *m* + *n* = *N*. In second-quantized language, the bulk part is given by

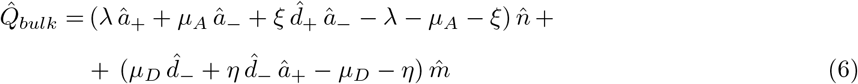

and the edge correction is given by

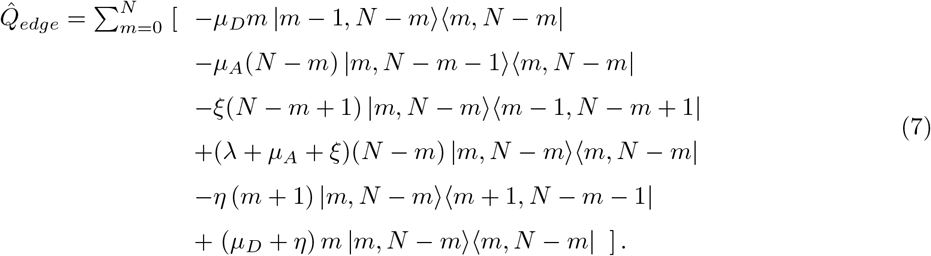

In Eq. (7), 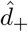 and 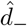 are respectively creation and annihilation operators for dormant tumor foci and 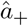 and 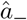 are those for the active foci. These operators are defined as:

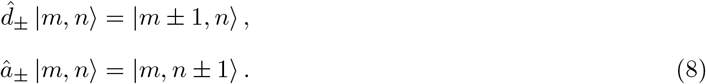

The number operators 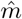 and 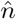 are defined in the usual fashion:

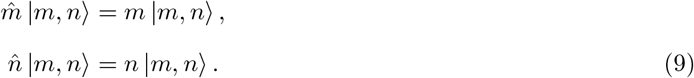

The structure of the matrix 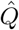 is block-tridiagonal in the linear space 𝒟 ⊗ 𝒜, with block indices (*m, m*^1^) that run across the Fock states of 𝒟, and with indices (*n, n^1^*) within each nonzero block that run across the Fock states of 𝒜. The *N* + 1 blocks along the diagonal are themselves tridiagonal and decrease in size as the 𝒟-space index *n* increases. The structure of 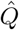, as described in Appendix A, is sufficiently complicated that explicit analytical solutions are not straightforward, although analytical formulas are available for the inversion of general tridiagonal and even certain types of block-tridiagonal matrices [31, 32]. Analytical and even stable numerical methods for general level-dependent QBD processes, i.e., QBD models with a block-tridiagonal matrix structure where the blocks are not constant along the diagonals [33], are scarce in the literature. Although a matrix-analytic method has been developed for these models in [34], it still relies on the ability to to solve non-trivial matrix equations. Numerical methods have been developed for finding stationary distributions in level-dependent QBD models [35], but generally applicable numerical methods for finding transient solutions (other than the expensive matrix exponentiation) have yet to be developed [36]. For a special class of level-dependent QBD models with applications in biology and epidemiology, a method based on a continuous-fraction representation of the Laplace-transformed transition probabilities has recently been developed [36]. However, it is not applicable to the model defined by Eqs. (5), (6) and (7), where all the transition and birth/death rates can be nonzero.

If we disregard constraint (1) for the moment and consider the special case *η* = *ξ* = *µ*_*D*_ = 0, the system reduces to a continuous-time birth-and-death process with transition rates *λn* and *µ*_*A*_*n* (otherwise known in queuing theory as the *M/M/*1*/N* queue [29]) and with absorbing boundary states. The version of this model with *N → ∞* (i.e., semi-infinite Markov chain) and reflecting boundary conditions has been studied extensively and analytical solutions have been obtained for its transient analysis using several techniques [37–41]. A version of the *M/M/*1*/N* queuing model that is more relevant to our analysis is the one with finite *N* and absorbing boundary states, which is solved analytically in [42], where the transient solution is obtained and a simple expression is given for the large-time probability *π*_*rec*_ of absorption at the state *n* = *N* (corresponding to recurrence in our model), under the initial condition *p*_*n*_(0) = *δ*_*n,n*0_:

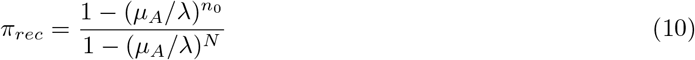

The probability of absorption at the zero-particle state *n* = 0 (corresponding to cure) is then *π*_*cure*_ = 1 *− π*_*rec*_. In the limit *N* → *∞*, we note that this model has a phase transition at *µ*_*A*_/*λ* = 1:

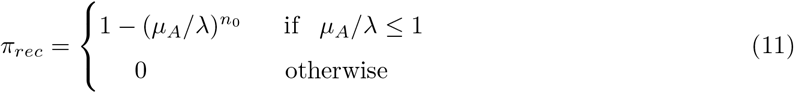

As will be shown in Section 5.1, a similar stationary solution also occurs in general in the QBD model defined by Eqs. (5), (6) and (7). We will show this both analytically in the continuum limit and in simulations of the discrete-state stochastic process.

## 4. Continuum-limit of the discrete-state QBD model

A simple approach that is suitable for our tumor recurrence model is to take the continuum limit of the master equation (5), i.e., take the large *N* limit. Since the reciprocal of the detectable tumor size (1*/N*) is a natural small parameter in the model, the master equation can be expanded in powers of 1*/N* and thus converted to a continuum equation. The resulting partial differential equation may then be solved using well-developed methods [43].

There are two alternative ways to represent the time evolution of the stochastic process at hand [43–47], namely the *forward* master equation (5) and the *backward* master equation

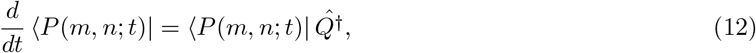

where 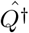 is the adjoint of the operator 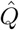 defined in Eqs. (6), (7) and the state vector *〈P*(*m, n*; *t*)*|* is defined as the probability to end up at a particular state 〈*m, n*| at time *t*, starting from *any* state *(m*_0_, *n*_0_| at time *t* = 0, i.e.,

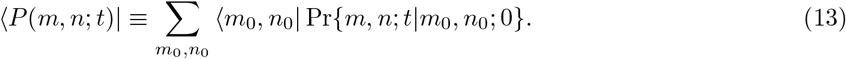

When either Eq. (5) or Eq. (12) is used in the large-*N* expansion, we get respectively the *forward* or the *backward* Kolmogorov equation in the continuum limit 1*/N «* 1 by truncating the expansion after the second term, as discussed further below.

### 4.1 Forward Kolmogorov approach

We can define a continuum limit of the state space Ω = *{*(*m, n*)*|m ≥* 0; *n ≥* 0; *m* + *n ≤ N}* by defining continuous variables *x* = *m/N* and *y* = *n/N*, which represent the dormant and active foci populations as a fraction of the detectable tumor size *N*, respectively. We define a “tumor focus” as the resolution scale of our model: for example, we can define a focus as 1*/*1000 of a detectable tumor, in which case *N* = 1000 is a natural definition of recurrence (detectable tumor size). When *N »* 1, the lattice of discrete states (*m, n*) becomes a continuum as the spacings 1*/N* decrease to zero. The discrete probabilities *p_mn_*(*t*) can then be replaced by a smooth probability density 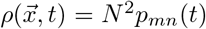, where 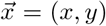 is an arbitrary point in the bounded region Ω = *{*(*x, y*)*|x ≥* 0; *y ≥* 0; *x* + *y ≤* 1*}*.

We proceed to take the continuum limit of the master equation by first replacing the raising/lowering operators (see Eqs. (5), (6) and (8)) by the corresponding translation operators in the continuum, i.e., 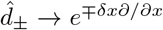 and 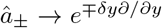 (the reason for the opposite signs is that the creation and annihilation operators are passive transformations, whereas the continuum translation operators are defined as active transformations). Then a Kramers-Moyal expansion of the master equation can be obtained by expanding the operator (6) in powers of *δx* = *δy* = 1*/N*. Retaining only the first and the second terms in this large-size expansion, we obtain the two-dimensional Fokker-Planck equation

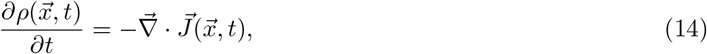

which is a local continuity equation with a probability current density given by

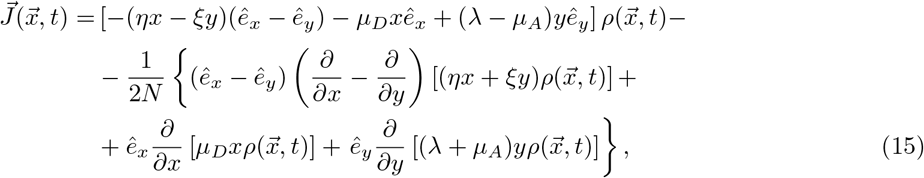

where each drift term is a product of the respective transition rate by the probability density, along the unit vector in the direction of the transition, and the terms proportional to 1*/N* represent diffusion, with a diffusion tensor that involves off-diagonal terms (i.e., the terms that involve mixed second derivatives *∂*^2^*ρ/∂x∂y* are nonzero) and is dependent upon the state variables (*x, y*).

Since Eq. (14) gives the probability density 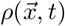 at any state 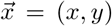 at time *t ≥* 0, given the initial condition 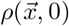, it corresponds to the well-known *forward* Kolmogorov equation [43–47]. Here we define the initial condition to be sharply peaked at a state 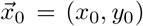, i.e., 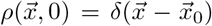. Note that the normalization of the probability density 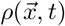 is *not* preserved by the time evolution, because the probability flux exits through the recurrence boundary *x* + *y* = 1, and also accumulates at the cure state (0, 0).

The boundary conditions for Eq. (14) are the following. At the recurrence line *x* + *y* = 1, the absorbing boundary condition is 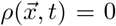. In the vicinity of the cure state (0, 0) we define 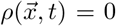 on the line *x* + *y* = *ε*, where *ε «* 1*/N* is a small parameter. In the weak limit (in the distributional sense) *ε →* 0, the small region *x* + *y ≤ ε*, representing the cure state, becomes a single absorbing point where the probability density collapses to a Dirac delta function weighted by the probability of cure before time *t*, which will henceforth be denoted by *p*_*cure*_(*t*).

On the boundaries *x* = 0, *ε ≤ y ≤* 1 and *y* = 0, *ε ≤ x ≤* 1, the boundary conditions are both reflecting, i.e., the normal component of the probability current density must vanish. In other words, 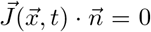, where 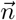 is the outward normal and the current density 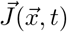 is defined by Eq. (15) (see Appendix B.1 for details).

Equation (14) is separable with respect to time, i.e., it can be reduced to an eigenvalue problem for the *forward* Fokker-Planck operator 𝓛_*f*_ defined by recasting Eq. (14) in the form 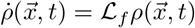. However, it is not separable in the coordinates (*x, y*) and also depends on these variables explicitly through coefficients *inside* the differential operators. Finding a basis of two-variable eigenfunctions of 𝓛_*f*_ satisfying the mixed boundary conditions described above is a difficult problem, but unnecessary for our main goal, which is to derive an equation for the *recurrence-free survival function S*(*t*), defined as the probability of no-recurrence before time *t*, i.e.,

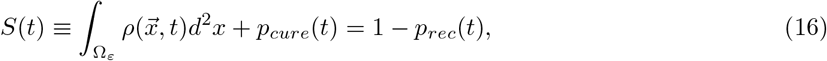

where the domain of integration is the region Ω_*ε*_ = *{*(*x, y*)*|x ≥* 0; *y ≥* 0; *ε ≤ x* + *y ≤* 1*}* and *p*_*rec*_(*t*) and *p*_*cure*_(*t*) are respectively the time-dependent probabilities of recurrence and cure before time *t*.

The function *S*(*t*) defined in Eq. (16) is a bridge that connects data (survival curves) to the model. The *backward* Kolmogorov approach is often more appropriate to first-passage time problems [45–48] and will be used in combination with Eq. (15) to derive an equation for *S*(*t*). The current density derived from the forward equation (Eq. (15)) will be used to derive formulas for the probability flux into the cure state or through the recurrence boundary (note that unlike the forward equation, the backward equation cannot be expressed as a local continuity equation for the conservation of probability). Furthermore, the boundary conditions for the backward equation are derived from those of the forward equation, as explained in Appendix B.2.

### 4.2 Backward Kolmogorov approach

Starting with Eq. (12) and using the adjoint of the master equation operator (6), we can derive the backward Kolmogorov equation by means of a Kramers-Moyal expansion similar to that leading to the forward equation (14). It should be noted that the differential operators in the backward equation act on functions of the initial-state variables (*x*_0_*, y*_0_):

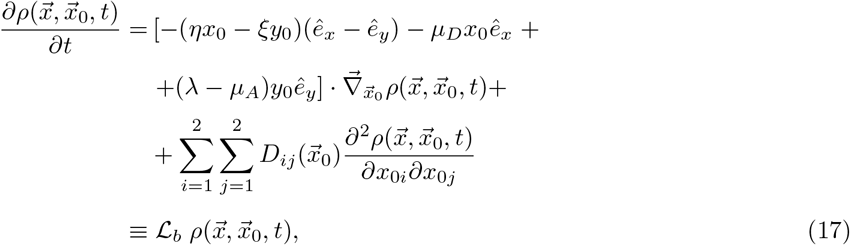

where 𝓛_*b*_ is the backward operator, (*x*_01_*, x*_02_) *≡* (*x*_0_*, y*_0_), and 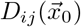 are the components of the diffusion tensor

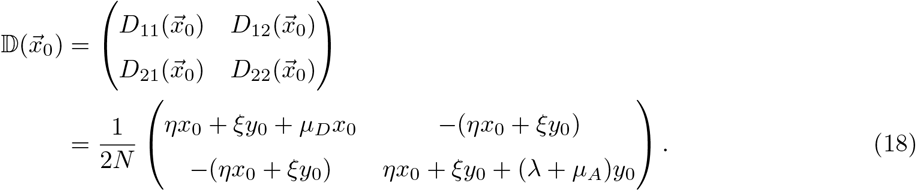

The backward equation (17) is somewhat different from Eq. (14) in that the non-constant coefficients appear *outside* the differential operators. It is subject to the *final* condition that some state 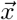 will be reached at time *t*, starting from *anywhere* 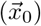 in the state space. This explains why the backward equation is often more useful for first-passage time problems than its forward counterpart.

The boundary conditions for the backward equation (17), which can be derived from those of the forward equation (14), are the following. On both absorbing boundaries *x*_0_ + *y*_0_ = *ε* and *x*_0_ + *y*_0_ = 1, 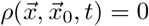. At the reflecting boundaries, the boundary conditions are

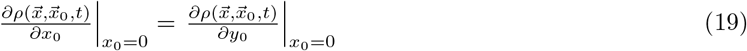

at the boundary *x*_0_ = 0, *ε ≤ y*_0_ *≤* 1 and

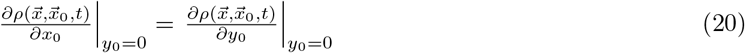

at the boundary *y*_0_ = 0, *ε ≤ x*_0_ *≤* 1. These boundary conditions are derived in Appendix B.2.

The equations for the probability flux through the absorbing boundaries *x*+*y* = 1 and *x*+*y* = *ε* are given in Appendix C. From them, we can derive the partial differential equations below for the time-dependent probabilities of recurrence and cure before time *t*, respectively (see Appendix C for details):

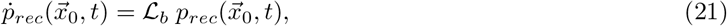

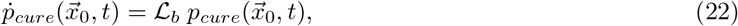

where 𝓛_*b*_ is the backward operator defined in Eq. (17) and we have used the notation 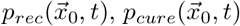 to recall the dependence on the initial condition 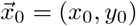. Since the recurrence-free survival function is defined as 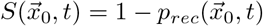, it must satisfy the PDE

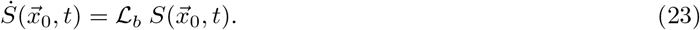

For *µ*_*D*_ = 0, we can simplify the backward equation (17) by transforming to the new variables *z*_0_ *≡ x*_0_, *w*_0_ *≡ x*_0_ + *y*_0_,

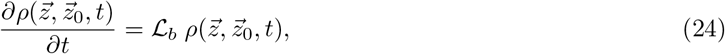

where 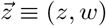 and the transformed backward operator is given by

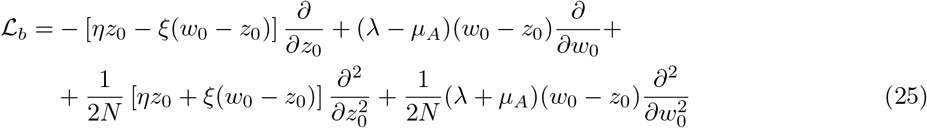

In the new variables (*z*_0_*, w*_0_), the boundary conditions for Eqs. (21), (22), (23) and (24) are summarized in Table 1. The Neumann boundary conditions for Eq. (24) at the boundaries *z*_0_ = 0 and *z*_0_ = *w*_0_, given in Table 1, follow immediately from Eqs. (19) and (20), and those for Eqs. (21), (22) and (23) follow from the equations for the probability flux through the absorbing boundaries *x* + *y* = 1 and *x* + *y* = *ε*, given in Appendix C.

**Table 1:** Boundary conditions for the PDEs satisfied by a few relevant functions of the initial-state variables 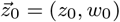, where *z*_0_ *≡ x*_0_ and *w*_0_ *≡ x*_0_ + *y*_0_.

| Function | Equation | Boundary |  |  |  |
| --- | --- | --- | --- | --- | --- |
| | | $w_0 = \varepsilon$ | $w_0 = 1$ | $z_0 = 0$ | $z_0 = w_0$ |
| $\rho(\vec{z}, \vec{z}_0, t)$ | $\dot{\rho} = \mathcal{L}_b \rho$ | $\rho = 0$ | $\rho = 0$ | $\frac{\partial \rho}{\partial z_0} = 0$ | $\frac{\partial \rho}{\partial z_0} = 0$ |
| $p_{rec}(\vec{z}_0, t)$ | $\dot{p}_{rec} = \mathcal{L}_b p_{rec}$ | $p_{rec} = 0$ | $p_{rec} = 1$ | $\frac{\partial p_{rec}}{\partial z_0} = 0$ | $\frac{\partial p_{rec}}{\partial z_0} = 0$ |
| $p_{cure}(\vec{z}_0, t)$ | $\dot{p}_{cure} = \mathcal{L}_b p_{cure}$ | $p_{cure} = 1$ | $p_{cure} = 0$ | $\frac{\partial p_{cure}}{\partial z_0} = 0$ | $\frac{\partial p_{cure}}{\partial z_0} = 0$ |
| $S(\vec{z}_0, t)$ | $\dot{S} = \mathcal{L}_b S$ | $S = 1$ | $S = 0$ | $\frac{\partial S}{\partial z_0} = 0$ | $\frac{\partial S}{\partial z_0} = 0$ |
| $T_{rec}^{(1)}(\vec{z}_0)$ | $\mathcal{L}_b [p_{rec}(\infty) T_{rec}^{(1)}] = -p_{rec}(\infty)$ | undef.* | $T_{rec}^{(1)} = 0$ | $\frac{\partial T_{rec}^{(1)}}{\partial z_0} = 0$ | $\frac{\partial T_{rec}^{(1)}}{\partial z_0} = 0$ |

\* The mean recurrence time diverges at the cure-state boundary $w_0 = \varepsilon$ .

The PDEs (21), (22) and (23), subject to the boundary conditions given in Table 1, are all separable in time, but not in the initial-state variables (*z*_0_*, w*_0_). However, they can be solved analytically in the large-time limit *t → ∞*, as will be shown in Section 5.1 below.

A key random variable in our model is the recurrence time (which will be henceforth denoted by *T*_*rec*_), corresponding to the first passage time through the recurrence boundary. The normalized probability that *T*_*rec*_ lies between *t* and *t* + *dt* can be obtained from a simple application of Bayes’ theorem [47]:

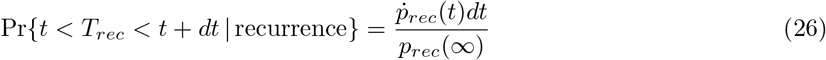

In other words, the probability density function (PDF) for the recurrence time *T*_*rec*_ is 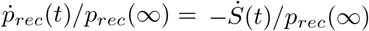, where the probability 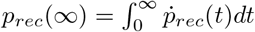 that recurrence takes place at any time *t < ∞* is given in closed analytic form in Section 5.1 below (see Eq. (32)). The ratio *p*_*rec*_(*t*)*/p*_*rec*_(*∞*) then gives the cumulative distribution function (CDF) of *T*_*rec*_.

Let

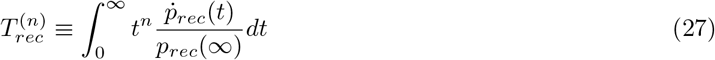

denote the *n*-th moment of the recurrence time *T*_*rec*_ (*n* = 0, 1, 2*, …*). From Eqs. (21) and (27), we get the hierarchy of equations below [44, 45], where the *n*-th moment of *T_rec_* is related to its (*n −* 1)-th moment:

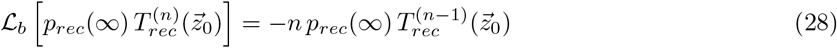

Here we have used the notation 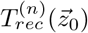 to recall the dependence on the initial condition 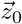. Since the function *p*_*rec*_(*∞*) also depends on the initial condition through the variable *w*_0_ (see Eq. (32) below), it cannot be taken outside the backward operator 𝓛_*b*_, because the latter acts on the initial condition variables (*z*_0_*, w*_0_).

The boundary condition for Eq. (28) at the absorbing boundary *w* = 1 is 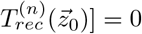. From Eq. (27) and Table 1, it also follows that Neumann boundary conditions 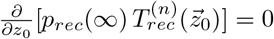 must be imposed at both reflecting boundaries *z*_0_ = 0 and *z*_0_ = *w*_0_.

Letting *n* = 1 in Eq. (28), we get an equation for the mean recurrence time (MRT), denoted here by 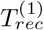:

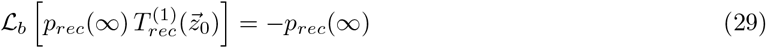

This is a key equation in our analysis, which will be used to find an approximate analytical formula for the MRT.

## 5. Results and discussion

In this section, we discuss our main results, namely the stationary solution of Eq. (21) at large times in closed analytic form, as well as the “outer solution” of the mean recurrence time (MRT) equation (29) at leading (zeroth) order in 1*/N*, which is approximately valid everywhere except inside thin boundary layers that stretch along the reflecting barriers *z*_0_ = 0 and *z*_0_ = *w*_0_. For several choices of the parameters and initial conditions, these solutions are compared against simulations. We also describe a simple procedure to fit the model to survival data, using serous ovarian cancer data downloaded from the public database *The Cancer Genome Atlas* [54] as an example.

### 5.1 Stationary solution at large times

The stationary solution of Eq. (21), which satisfies 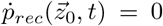, gives the large-time probability *p*_*rec*_(*z*_0_, *w*_0_*, ∞*) of absorption at the recurrence boundary *w* = 1, given the initial state (*z*_0_*, w*_0_). In the large-time limit *t → ∞*, the probability of cure is given by 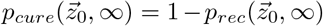: recurrence or cure are the only possible fates at large times. Therefore, *p*_*rec*_(*z*_0_*, w*_0_*, ∞*) satisfies the *homogeneous* backward equation

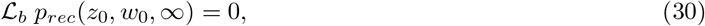

where the backward operator in the variables (*z*_0_*, w*_0_) is given by Eq. (25). The boundary conditions are given in Table 1.

An ansatz solution to Eq. (30) would be a recurrence probability that only depends on the *total* initial number of tumor foci *w*_0_ and not specifically on what fraction of this initial number are dormant foci (*z*_0_), i.e., 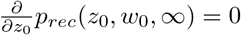. This type of solution automatically satisfies the Neumann boundary conditions at the reflecting boundaries *z*_0_ = 0 and *z*_0_ = *w*_0_ (see Table 1). Using the ansatz above and Eq. (25), the homogeneous PDE (30) becomes the following ODE in the variable *w*_0_:

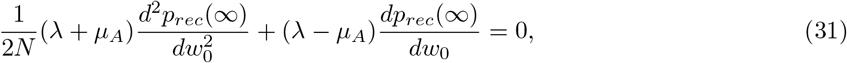

where *p*_*rec*_(*∞*) *≡ p*_*rec*_(*z*_0_*, w*_0_*, ∞*) only depends on *w*_0_. This equation can be easily solved for the Dirichlet boundary conditions given in Table 1. The solution is

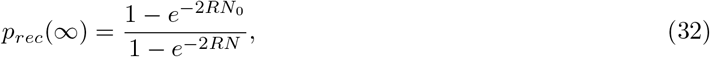

where *N*_0_ = *w*_0_*N* is the initial total number of tumor foci and

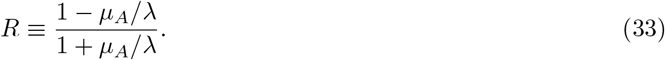

In Eq. (32), the limit *ε →* 0 has been taken. In the large-time limit *t → ∞*, the probability of cure is given by

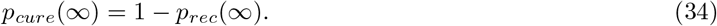

In Fig. 2, the function (32) is plotted for different values of the detectable tumor size *N* at a fixed initial number of foci *N*_0_ and also for different initial conditions *N*_0_ at a fixed *N*.

**Figure 2:**
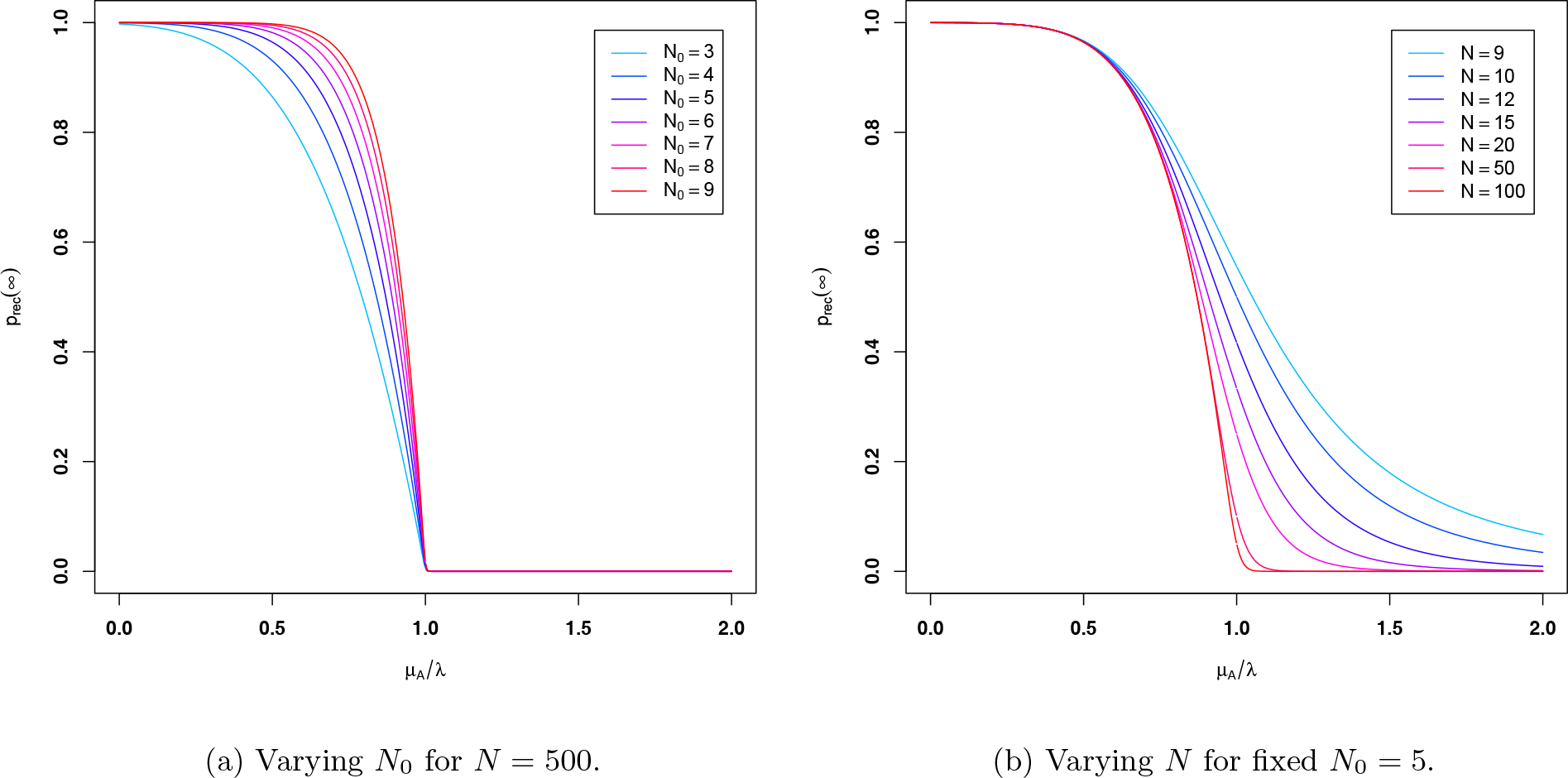
Large-time probability of recurrence *vs. µ*_*A*_/*λ* for *µ*_*D*_ = 0, obtained analytically using the backward Kolmogorov approach (see Eq. (32)). Pane 2a shows the effect of changing the total initial number of tumor foci (*N*_0_), whereas pane 2b shows the effect of a finite detectable-tumor size *N* on the phase transition.

In the limit *N → ∞*, this solution displays a phase transition at *µ_A_/λ* = 1:

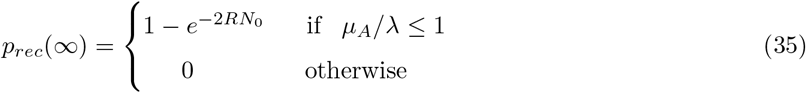

Thus, in the limit *N → ∞*, the large-time probability of cure for *µ*_*A*_/*λ* ≤ 1 is *p*_*cure*_(*∞*) = 1 − *p*_*rec*_(*∞*) = *e*^−2*RN*0^, which is the large-time limit of the recurrence-free survival function *S*(*t*).

The drift term of the Fokker-Planck equation (14) alone is not sufficient to reproduce the phase transition (35). The latter is the result of a combination of drift *and* diffusion in the presence of two opposite absorbing boundaries, so that the initial delta peak 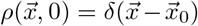 splits into a bi-modal probability density 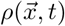 with two peaks that travel in opposite directions, toward the cure or recurrence boundaries. For *µ*_*A*_/*λ* > 1, the height of the peak traveling toward the recurrence boundary vanishes, so the final state in the large-time limit *t* → *∞* becomes the cure state (0, 0), with probability 1. The result (32) is only slightly different from the stationary solution of the M/M/1/N queue with absorbing boundary states (see Eq. (11)). As we will show, it agrees with simulations of the discrete-state QBD process.

### 5.2 Approximate solution of the mean recurrence time equation

In this section, we will solve the mean recurence time equation (29) analytically to the leading (zeroth) order in 1*/N*. This approximation is valid outside boundary layers near the reflecting barriers *z*_0_ = 0 and *z*_0_ = *w*_0_, the sizes of which vanish as *N → ∞*. The leading order solution can be obtained by neglecting the second derivative terms in Eq. (29). This results in the first order PDE

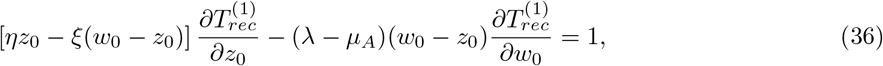

which can be solved by the method of characteristics. The solution is

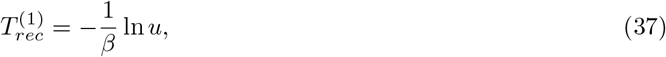

where *u* is the only root of the transcendental equation

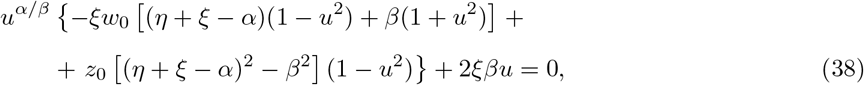

where

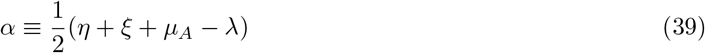

and

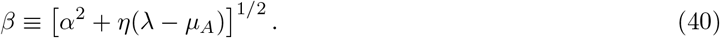

The procedure for obtaining this solution is described in Appendix D. The finite-*N* solution of Eq. (29) converges to the leading-order approximation given by Eqs. (37) and (38) pointwise, but not uniformly. Indeed, the approximate solution above does not satisfy the homogeneous Neumann boundary conditions at *z*_0_ = 0 or *z*_0_ = *w*_0_ (see Table 1); near each reflecting barrier, there is a boundary layer within which the zeroth-order approximation in 1*/N* fails. This is seen in the form of the mean recurrence time (MRT) level curves given by Eqs. (37) and (38), which are straight lines that are not parallel to the recurrence boundary *w*_0_ = 1, except asymptotically in the limit 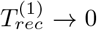 (see Fig. 3). In reality, close to each reflecting barrier (within some distance that vanishes in the limit *N → ∞*), the exact (finite-*N*) level curves bend toward the boundary, where these curves end at a right angle. At fixed initial conditions and as a function of the parameters *µ*_*A*_/*λ* and *ξ*/*η*, the shapes of the MRT level curves given by Eqs. (37) and (38) in parameter space are shown in Fig. 4 for two different initial conditions, namely *N*_0_ = 7, *m*_0_ = 3 and *N*_0_ = 50, *m*_0_ = 20.

**Figure 3:**
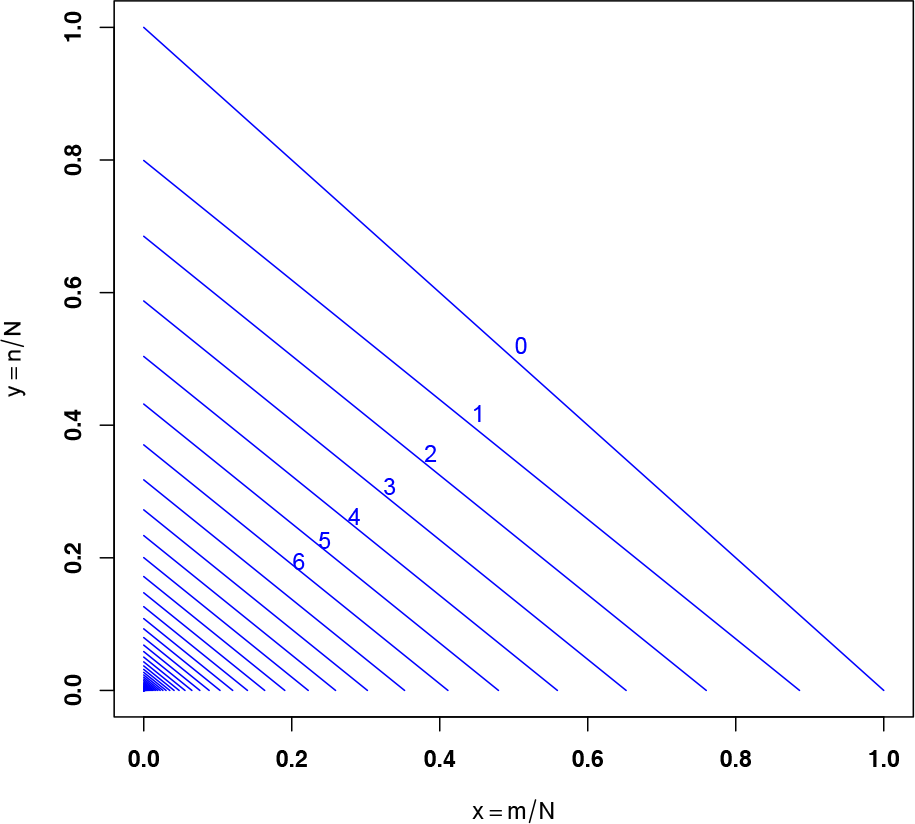
Level curves of the leading-order outer solution of the mean recurrence time equation (see Eqs. (37) and (38)) for *µ*_*A*_/*λ* = 0.5 and *ν* ≡ *ξ*/*η* = 2.5. At large *N*, these curves are approximately valid outside boundary layers that exist close to the reflecting barriers *x*_0_ = 0 and *y*_0_ = 0. The sizes of these boundary layers vanish in the limit *N → ∞*. The values of 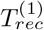 at the first few level curves are given in the figure in units of the doubling time 1*/λ*. The spacings between adjacent lines (which were plotted for 1*/λ* increments of 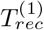 approach zero at the cure state (0, 0), where the mean recurrence time diverges logarithmically.

**Figure 4:**
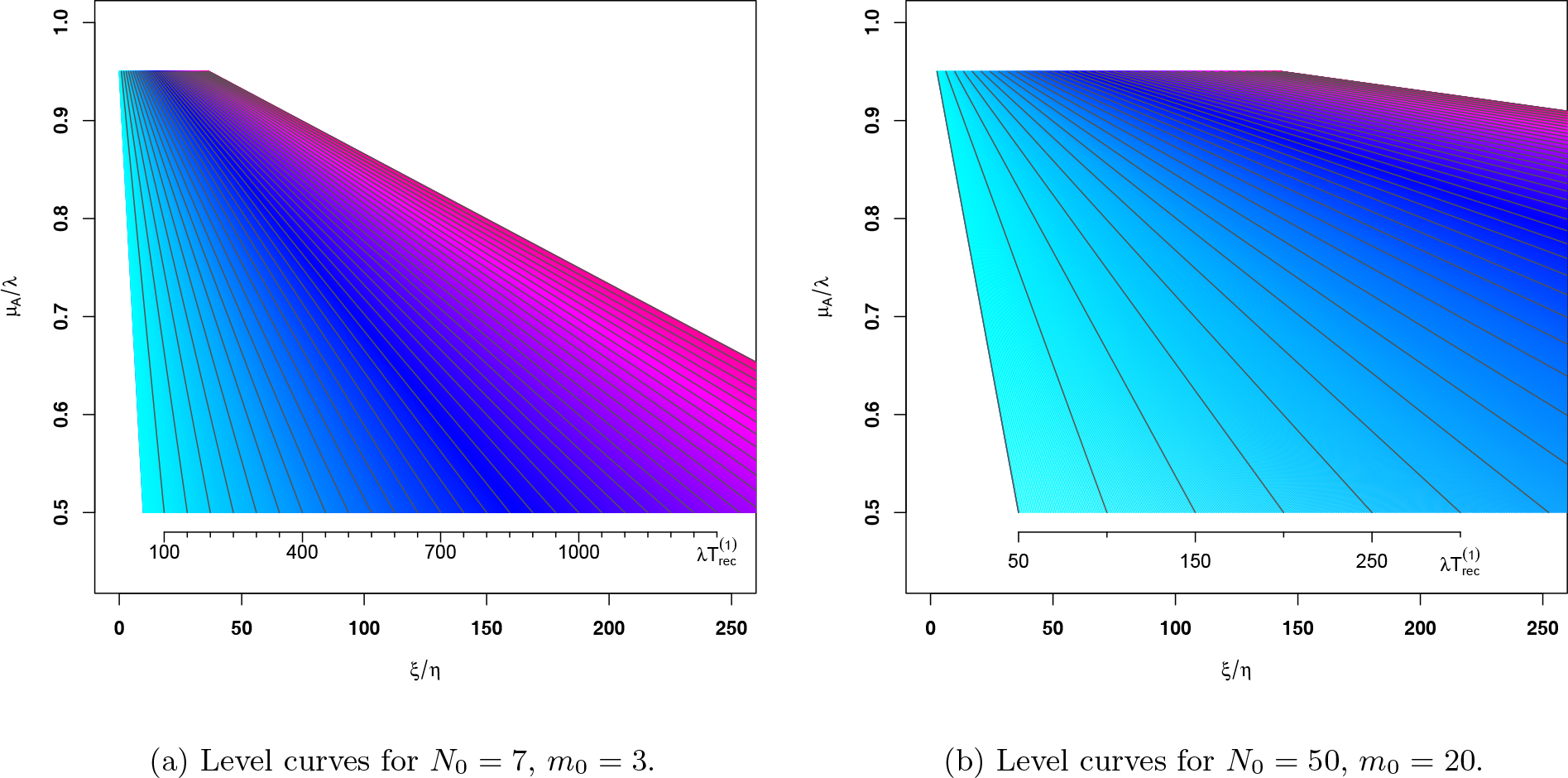
Level curves in parameter space of the leading-order outer solution of the mean recurrence time (MRT) equation (see Eqs. (37) and (38)) for two different initial conditions, with *N* = 100. The ruler at the bottom of each plot gives the values of the MRT in units of the doubling time 1*/λ*.

The PDE (29), along with its boundary conditions given in Table 1, is a singular perturbation problem that should be handled by special perturbation methods, because the small parameter 1*/N* premultiplies the second-derivative terms in the backward operator (25). Further inspection shows that Eq. (29) is structured in such a way that the problem can be treated by boundary-layer theory [49]. Since the solution of Eq. (29) obtained by dropping the diffusion (1*/N*) terms in the backward operator (25) is only valid *outside* the boundary layers that exist near the reflecting boundaries *z*_0_ = 0 and *z*_0_ = *w*_0_ (the sizes of which vanish as *N → ∞*), in the language of boundary layer theory the solution given by Eqs. (37) and (38) is called the “outer solution” of the boundary value problem. The “inner solutions”, on the other hand, require proper rescaling of the variables before these solutions can be expanded asymptotically in the parameter 1*/N*; in this case, the second-derivative terms in Eq. (25) cannot be neglected inside each boundary layer, since they become comparable to the first-derivative terms within each layer.

An approximate composite solution that would be valid everywhere can in principle be obtained by the method of matched asymptotic expansions, which requires solving Eq. (29) both inside and outside the boundary layers [49]. In this paper, however, only the leading-order outer solution is given (Eqs. (37) and (38) above). In practice, the outer solution itself is already a quite useful result, even at the lowest order in 1*/N*, since it agrees reasonably with simulations of the model (as will be shown in Section 5.3 below), except for small corrections that can in principle be calculated using the boundary-layer method.

### 5.3 Simulations

The discrete-state QBD process was simulated for several values of the model parameters and initial conditions using a pseudo random number generator. The results from the simulations were then compared to the large-time stationary solution given by Eq. (32), and also to outer solution given by Eqs. (37) and (38). The time step was chosen to be such that the transition probabilities would always remain sufficiently small within the range of the transition rates used in the simulations, even for transitions between states with large numbers *m, n ∼ N*. For a given set of parameters, each simulation tracked both the fraction of patients for which the tumor recurred and the recurrence times (from which the MRT was estimated) for an ensemble of 400 hypothetical patients. The detectable tumor size was set to *N* = 100.

For a simulation with *ν* ≡ *ξ*/*η* = 5 and initial condition *m*_0_ + *n*_0_ = 7, *m*_0_ = 3, Fig. 5 shows a plot of the large-time fraction of patients for which the tumor recurred against *µ*_*A*_/*λ*. The figure also shows that the result of the simulation agrees with the analytical formula given by Eq. (32). Fig. 6 shows a plot of the MRT obtained in simulations against the initial total number of tumor foci (*N*_0_) along the line *m*_0_ = *n*_0_ = *N*_0_*/*2, for 4 different values of *µ*_*A*_/*λ*, with *ν ≡ ξ/η* = 2.5. These results are compared to the smooth curves obtained from the leading-order outer solution of the MRT equation (Eqs. (37) and (38)). At the cure state *N*_0_ = 0, the MRT diverges logarithmically and the discrepancy between simulations and the outer solution increases as *N*_0_ approaches zero, due to the boundary-layer structure (the vicinity of the cure state is the region where the boundary layers along the reflecting barriers overlap).

**Figure 5:**
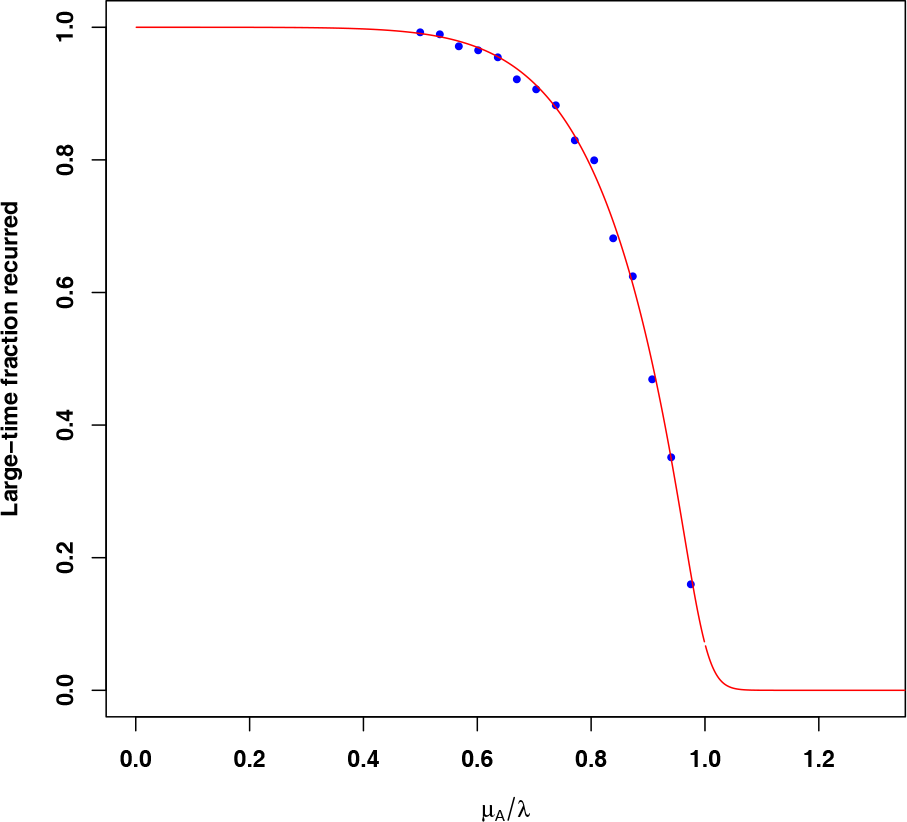
Large-time fraction of patients for which the tumor recurred, obtained in a simulation of the QBD model with 400 hypothetical patients. The parameters are *µ*_*D*_ = 0, *ν* ≡ *ξ*/*η* = 5, *m*_0_ + *n*_0_ = 7, *m*_0_ = 3 and *N* = 100. The red curve shows the analytical result given by Eq. (32).

**Figure 6:**
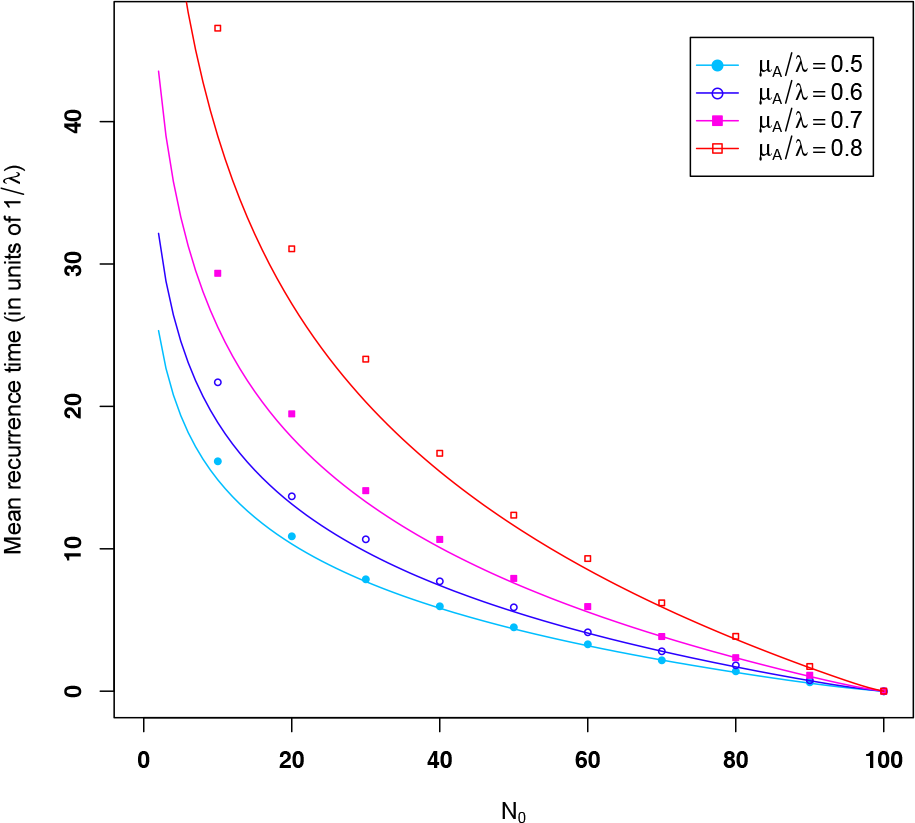
Mean recurrence time (MRT) obtained in simulations *vs.* initial total number *N*_0_ along the line *m*_0_ = *n*_0_ = *N*_0_*/*2, for different values of *µ*_*A*_/*λ* with *ν* ≡ *ξ*/*η* = 2.5 and *N* = 100. The simulations were run for 400 hypothetical patients. The smooth curves represent the leading-order outer solution of the MRT equation at different values of *µ*_*A*_/*λ* (see Eqs. (37) and (38)). We note the logarithmic divergence at the cure state *N*_0_ = 0.

For both initial conditions *N*_0_ = 7, *m*_0_ = 3 and *N*_0_ = 50, *m*_0_ = 20, Fig. 7 shows the MRT obtained in simulations against the ratio *ν ≡ ξ/η* at 5 different values of *µ*_*A*_/*λ*, and compares it to the curves obtained from the leading-order outer solution given by Eqs. (37) and (38). As expected, the higher the value of *µ*_*A*_/*λ*, the larger is the discrepancy, since the finite-size effect is greatest near the critical point *µ*_*A*_/*λ* = 1.

**Figure 7:**
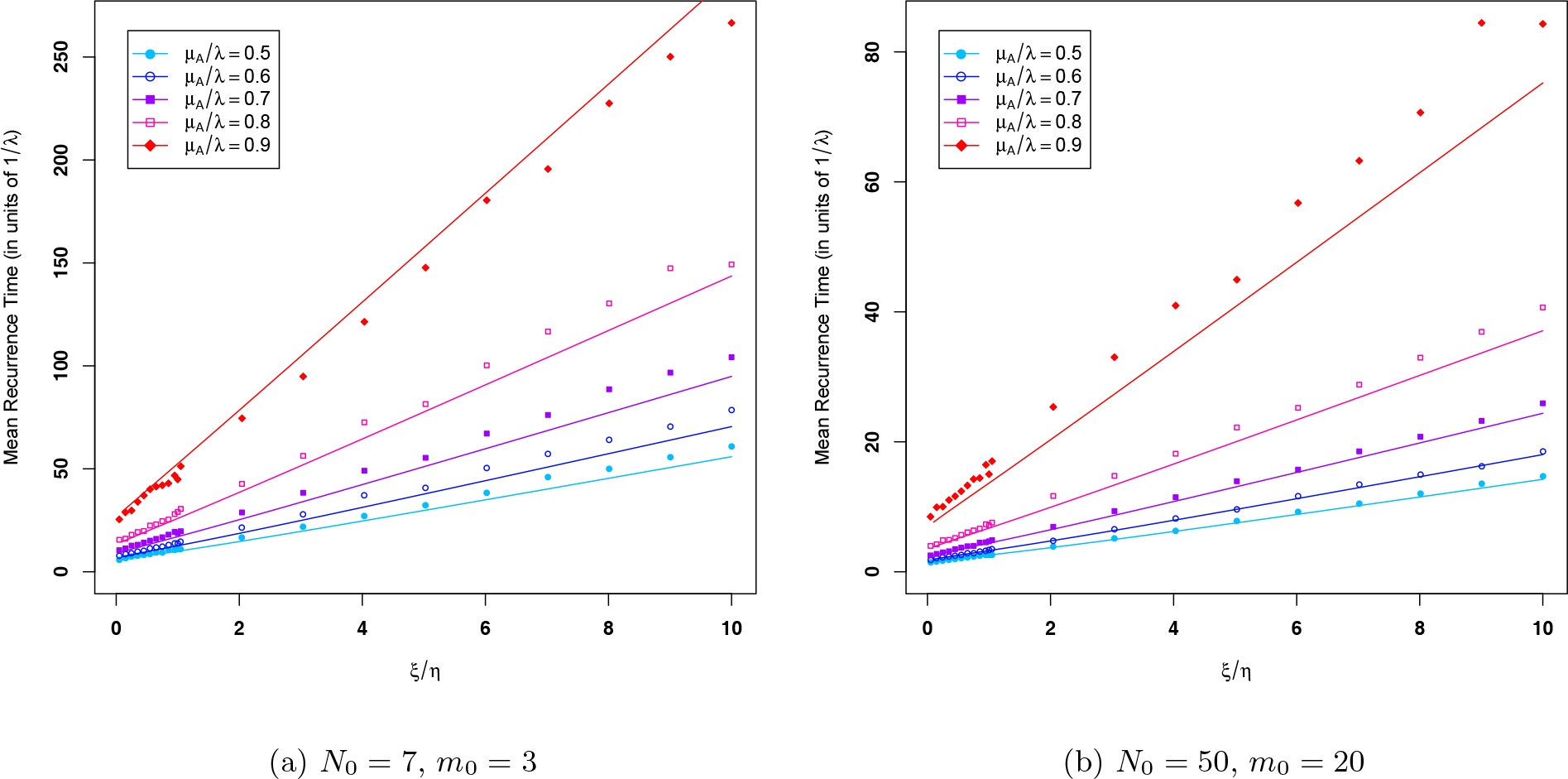
Mean recurrence time (MRT) obtained in simulations *vs.* the ratio *ν* ≡ *ξ*/*η* for two different initial conditions, with *N* = 100. Each set of points on each panel corresponds to a different value of *µ*_*A*_/*λ* and the lines represent the leading-order outer solution of the MRT equation (see Eqs. (37) and (38)). The results represent an average over 400 hypothetical patients.

It is worth noting that the ratio *ν ≡ ξ/η* can in principle be measured in experiments by estimating the relative times that the cells in a tumor spend in dormant versus active phases of the cell cycle, for example, through reconstruction of cell cycle dynamics from single-cell transcriptome data [50–52].

Fig. 8 shows the MRT obtained in simulations against *µ*_*A*_/*λ* for the initial conditions *N*_0_ = 7, *m*_0_ = 3 and *N*_0_ = 50, *m*_0_ = 20, at different values of the ratio *ν ≡ ξ/η*. These results are compared to the smooth curves obtained from Eqs. (37) and (38). At leading order, the MRT given by the outer solution diverges at the critical point *µ*_*A*_/*λ* = 1, as shown by Eqs. (37) and (38). The larger discrepancy near the critical point can be explained by a finite size effect in the phase transition given by Eq. (32), which effectively shifts the critical point to a value slightly higher than *µ*_*A*_/*λ* = 1 (see plot in Fig. 2b).

**Figure 8:**
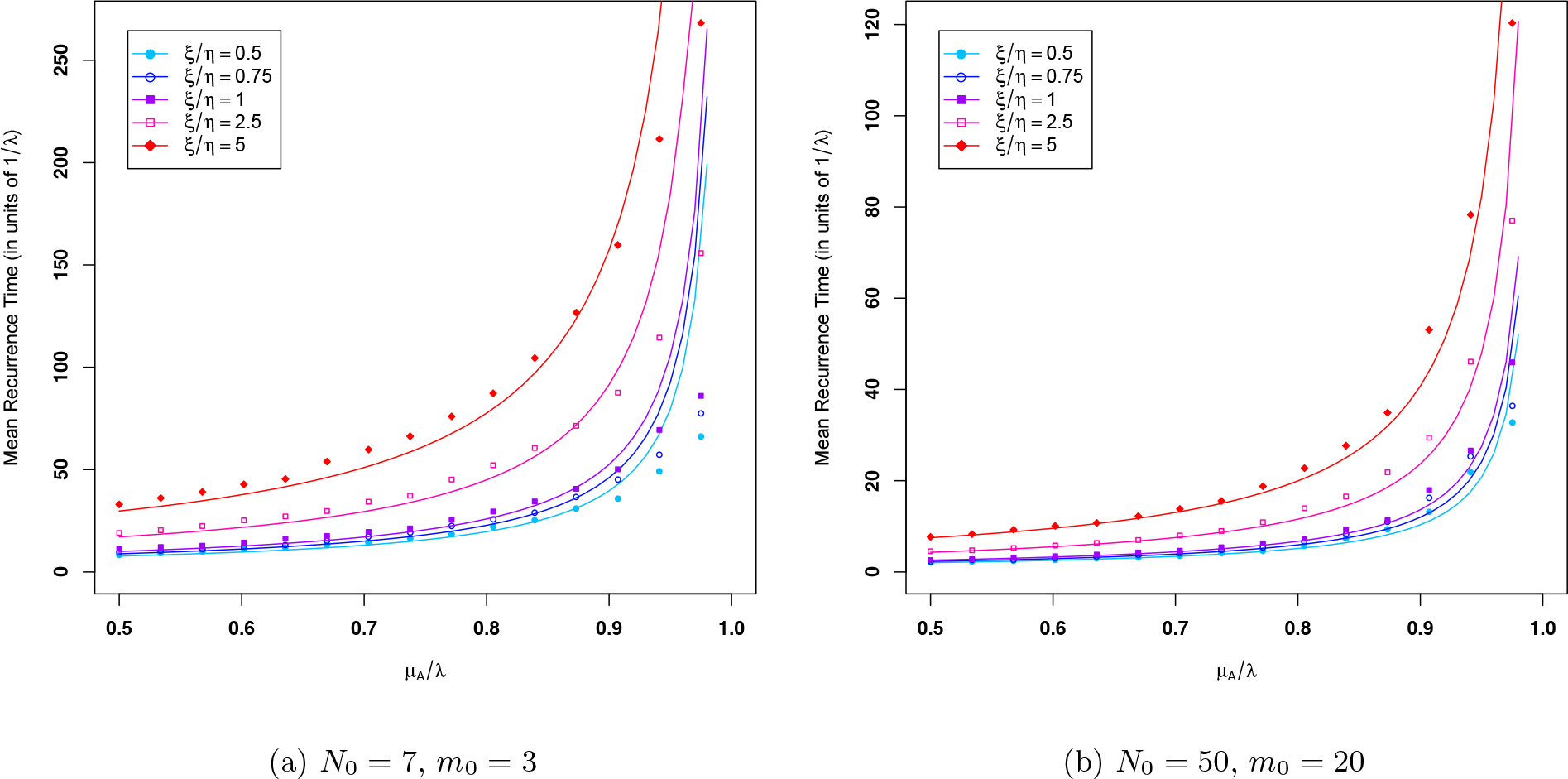
Mean recurrence time (MRT) obtained in simulations *vs. µ*_*A*_/*λ* for two different initial conditions, with *N* = 100. Each set of points on each panel corresponds to a different value of the ratio *ν* ≡ *ξ*/*η* and the smooth curves represent the leading-order outer solution of the MRT equation (see Eqs. (37) and (38)). The simulations were run for 400 hypothetical patients. At leading order, the MRT given by the outer solution diverges at the critical point *µ*_*A*_/*λ* = 1.

We note that the continuum limit of the discrete-state master equation is a good approximation, since it agrees reasonably well with the simulations even at leading (zeroth) order in 1*/N*. This is a remarkable feature of our model, since the diffusion limit sometimes fails for other types of birth-and-death processes, especially ones that involve non-linear transition rates [53].

### 5.4. Fitting the model to survival data

Even in the context of univariate birth-and-death processes, estimating model parameters from data is generally a difficult problem [55]. In this section, we describe a simple procedure to fit the model to survival data through an example. Since the method that we describe below relies on the leading-order outer solution given by Eqs. (37) and (38), it is approximately valid (within *O*(1*/N*) corrections) for initial conditions outside the boundary layers.

Serous ovarian cancer data downloaded from the public database *The Cancer Genome Atlas* [54] (TCGA) were used to generate the recurrence-free survival function shown in Fig. 9. Since 170 out of the 583 patients in the data set were censored, the Kaplan-Meier product-limit estimator [56] was used to estimate the time-dependent probability of no recurrence.

**Figure 9:**
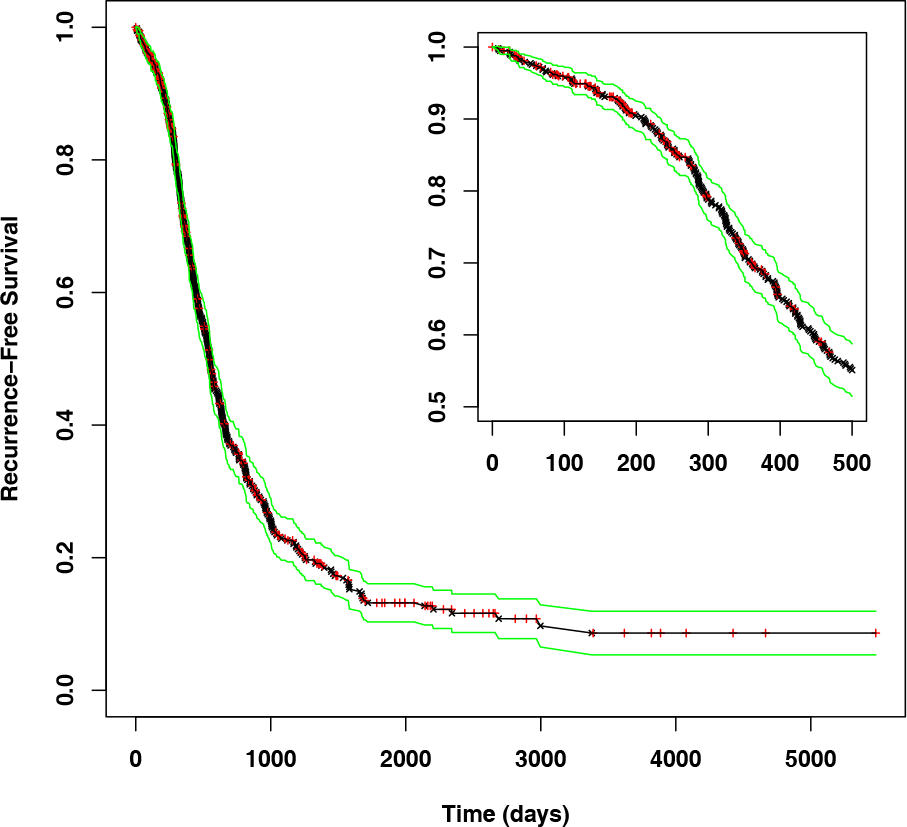
Product-limit estimate of the recurrence-free survival function (Kaplan-Meier curve [56]) for ovarian cancer in a group of 583 patients, obtained from TCGA data [54], showing Kaplan-Meier’s estimate for the 90% confidence interval (green lines). The red vertical crosses (+) represent censored patients, whereas the black saltire crosses (*×*) represent recurrence events. The inset provides a closer view of the KM curve for the time interval between 0 and 500 days.

In order to obtain the MRT from a given survival function *S*(*t*), we first need to renormalize the probability of recurrence *p*_*rec*_(*t*) = 1 − *S*(*t*) as in Eq. (26), i.e., the appropriate probability measure is *conditioned* on recurrence. The renormalized survival function 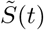 is then given by

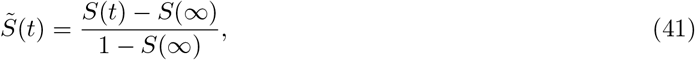

which vanishes in the limit *t* → *∞*. The MRT is then given by the area under the curve 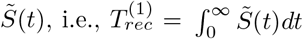. For the serous ovarian cancer data from TCGA, we found *S*(*∞*) = 0.086 (defined as the lowest value of the Kaplan-Meier estimate for *S*(*t*)) and an MRT of 687.5 days. For a given initial condition, estimation of the parameter *ν ≡ ξ/η* from Eqs. (37) and (38) requires that the time scale be fixed by specifying the doubling rate *λ*. Using the order of magnitude guess 1*/λ* = 40 days for the doubling time based on clinical data [57], the MRT for the ovarian cancer data in units of the doubling time was then fixed at *λ*(687.5 days) = 17.2.

For a given choice of the initial number *N*_0_, Eqs. (32) and (33) can be solved for the parameter *µ*_*A*_/*λ* using the value *p*_*rec*_(*∞*) = 1 *− S*(*∞*) = 0.914 obtained from the Kaplan-Meier curve. Assuming *m*_0_ = 0.4*N*_0_ (for which the outer solution gives a reasonable approximation of the MRT, unless *N*_0_ is too small), it then only remains to determine the value of the ratio *ν ≡ ξ/η* consistent with both the MRT (fixed at 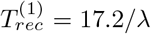) and the inferred *µ*_*A*_/*λ*. This is done by solving Eqs. (37) and (37) numerically for *ν* ≡ *ξ*/*η* (see also Eq. (2)).

Using the parameters determined through the scheme described above, survival curves were simulated for several initial conditions *N*_0_. In Fig. 10, these curves are shown in one plot, along with the Kaplan-Meier curve for the ovarian cancer data from TCGA. The time axis was rescaled to units of the MRT and the values of *N*_0_, *µ*_*A*_/*λ* and *ν ≡ ξ/η* used in the simulations are given in the legend. Among the initial conditions shown in the plot, the best fit corresponds to the choice *N*_0_ = 45.

**Figure 10:**
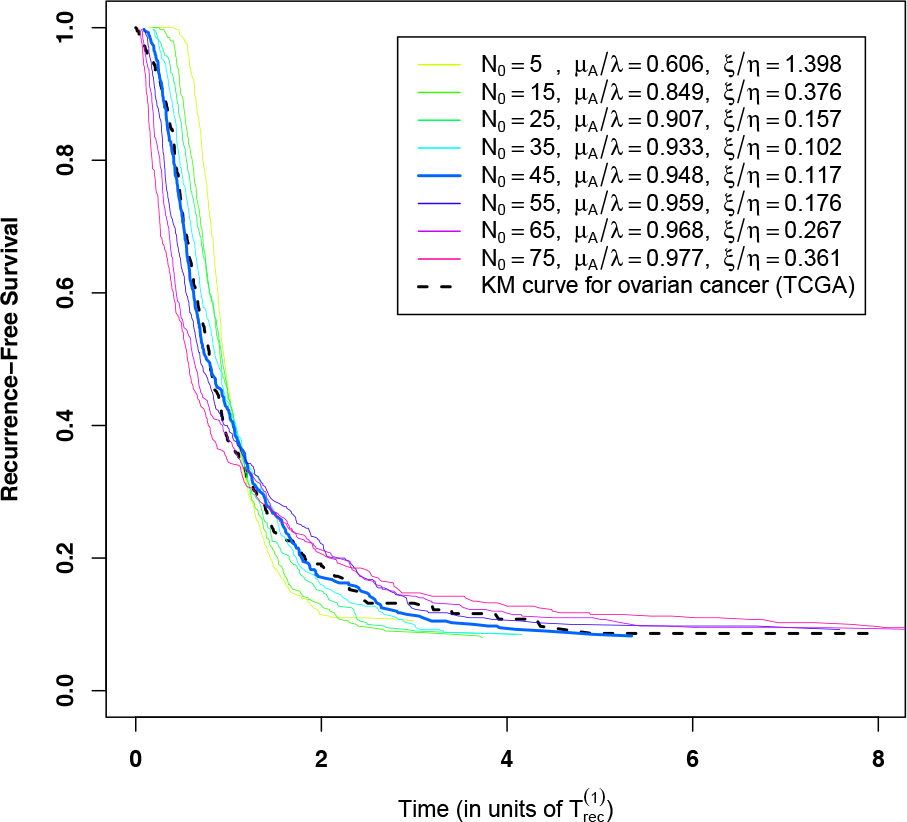
The same Kaplan-Meier curve shown in Fig. 9 for ovarian cancer data from TCGA [54] (dashed curve), plotted along with several recurrence-free survival curves obtained in simulations of the QBD model for 400 hypothetical patients, with *N* = 100. The mean recurrence time (MRT) for the TCGA data was found to be 687.5 days; assuming a doubling time of 1*/λ* = 40 days to fix the scale, the MRT in units of the doubling time was fixed at *λ*(687.5 days) = 17.2 for all the simulated curves. For each curve, the time axis was rescaled to units of the MRT. Each curve was simulated for a different value of *N*_0_ = *m*_0_ + *n*_0_ and assuming *m*_0_ = 0.4*N*_0_. For each value of *N*_0_, the parameters *µ*_*A*_/*λ* and *ν ≡ ξ/η* (given in the legend) were fixed using the scheme described in Section 5.4, which ensures that the MRT and the large-time recurrence probability both match the data. Among the initial conditions shown in the plot, the best fit (thick blue curve) corresponds to the choice *N*_0_ = 45.

In this procedure for fitting the model to data, we have assumed a sharply peaked initial condition, i.e., the initial condition is a delta function centered at some specified initial number of tumor foci *N*_0_. However, in reality this number should follow a probability distribution that would reflect the histogram of the residual-tumor size in the population under study. Moreover, different patients may have different responses to treatment (i.e., different values of *µ*_*A*_/*λ*), as well as different *ξ/η* values. This means that the fitted values should be regarded as only a guide that gives insight into the possible scenarios leading to the observed survival curve.

## 6. Conclusion

In this paper, we developed a mechanistic mathematical model aimed at describing the stochastic dynamics of tumor recurrence through a Quasi Birth-and-Death (QBD) process. The main assumption is the presence of residual tumor foci that can transition between a dormant, chemoresistant and an active-growth, chemosensitive state. We started with a continuous-time discrete-state master equation that describes the time-dependent probability *p*_*m,n*_(*t*) to be in a state with *m* dormant and *n* active tumor foci, and then showed that for a large detectable-tumor size *N*, the discrete master equation can be well approximated by a drift-diffusion equation in a continuous state space. Recurrence and cure were built into the model by imposing absorbing boundary conditions at the cure state (0, 0) and at the recurrence line *m* + *n* = *N*, respectively.

Using the forward and backward Kolmogorov approaches in the continuum limit, we derived an equation for the time-dependent probability of recurrence and the appropriate boundary conditions. The stationary solution at large times was then obtained analytically (see Eq. (32)) and we showed that it displays a phase transition as a function of *µ*_*A*_/*λ*, where *µ*_*A*_ is the death rate of active tumor foci and *λ* is their doubling rate. We also derived an equation for the mean recurrence time (MRT), which we solved analytically to leading order in 1*/N* by dropping the diffusion (second-derivative) terms in the equation, an approximation that works outside thin boundary layers along the reflecting barriers (see Eqs. (37) and (38) for the “outer solution”).

The analytical results were compared to simulations of the discrete-state QBD model. The large-time probability of recurrence obtained in simulations matched the analytical solution, whereas the MRT from the simulations showed a small discrepancy to the leading order outer solution of the MRT equation, except near the critical point *µ*_*A*_/*λ* = 1, where the discrepancy was larger due to the finite-size effect. In principle, it is possible to get an improved approximation by solving the MRT equation inside the boundary layer (where the variables have to be rescaled) and constructing a composite solution by the method of matched asymptotic expansions [49].

Finally, we described a scheme to fit the model to recurrence-free survival data (Kaplan-Meier curves), using ovarian cancer data from TCGA [54] as an example (Fig. 10). The model has potential applications to predicting the effect of changes in the tumor death rate or in the duration of chemotherapy on survival (recurrence rates). By switching the parameter *µ*_*A*_ at a specified time *t* = *t*_*chemo*_ (where *t*_*chemo*_ represents the duration of chemotherapy) from some specified level *µ*_*chemo*_ to a lower baseline level *µ*_0_, we can simulate the effect on survival of extending chemotherapy at lower dose (i.e., simultaneously increasing *t*_*chemo*_ and lowering *µ_chemo_*). This would allow quantitative studies of the effect of changes in chemotherapy regimens that can potentially be useful as a guide to clinical practice.

## 7. Acknowledgments

The authors would like to thank Alexandre Morozov for suggesting the continuum-limit approach, Siddhartha Sahi for a discussion on exact solutions of the discrete model, and Anshuman Panda for providing the data and for helpful discussions.

## Appendix A. Structure of the infinitesimal transition matrix

Projecting the operator 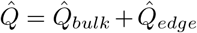 given by Eqs. (6) and (7) on both sides between basis vectors 〈*m, n*| and *|m′*, *n′*, we get matrix elements with a block-tridiagonal structure in the direct-product linear space 𝒟 ⊗ 𝒜:

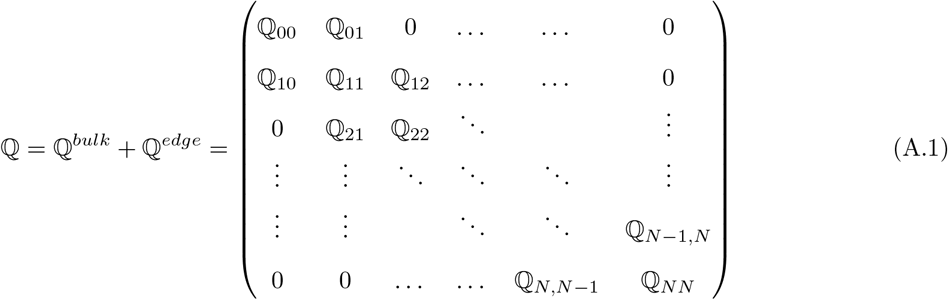

From the geometry of the state space boundary (see Fig. 1b), it follows that the block matrices 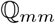 decrease in size as *m* increases: 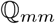 is an (*N − m* + 1) *×* (*N − m* + 1) matrix, since only the subspace spanned by the states *|m*, 0*), …, |m, N − m〉* is accessible. The bulk part of each block 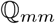 is tridiagonal and acts within the accessible subspace of 𝒜:

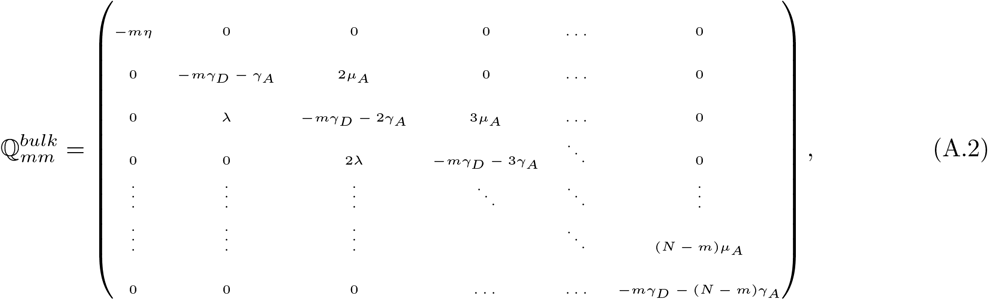

where

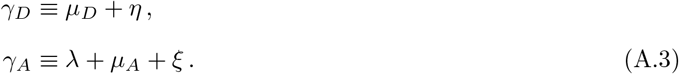

The bulk parts of the off-diagonal blocks are the matrices

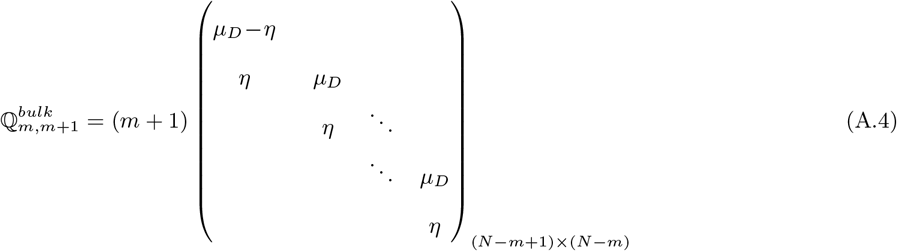

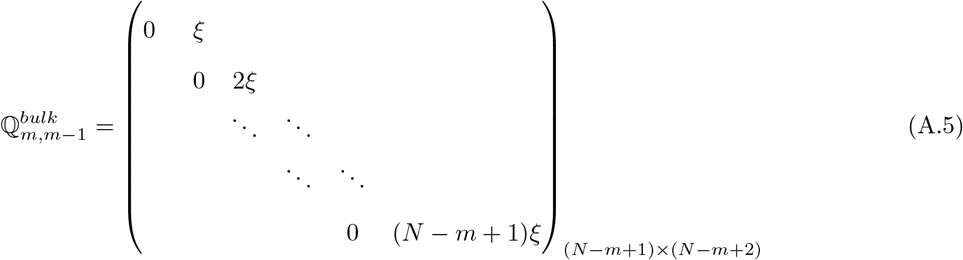

The edge corrections for the three block matrices above are given by

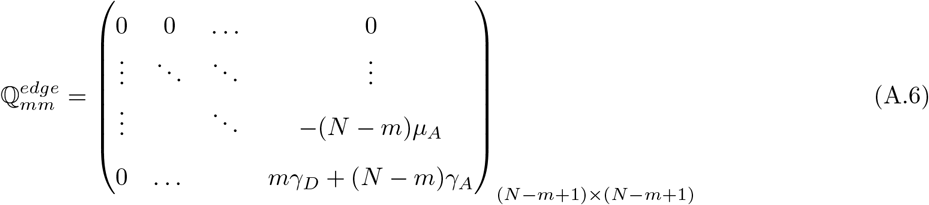

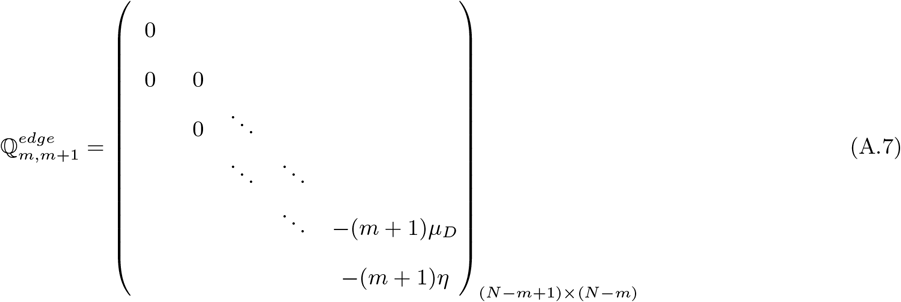

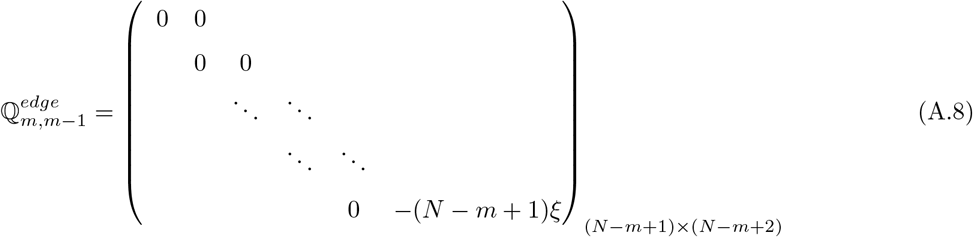

## Appendix B. Boundary conditions for the forward and backward Kolmogorov equations

In this appendix, we first explain in detail how the boundary conditions for the forward Kolmogorov equation are properly defined, with special note to the appropriate treatment of the single absorbing point at the origin (i.e., the cure state). Then we show how the boundary conditions for the backward Kolmogorov equation can be derived from those of the forward equation.

### Appendix B.1. Forward Kolmogorov equation

The boundary condition that the cure state 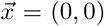 acts as a single absorbing point can be imposed by defining

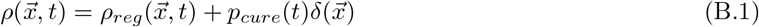

where 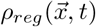 is the regular part of 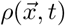 and *p*_*cure*_(*t*) is the probability of cure at any time *≤ t*. In Section 4.2, it is shown that this boundary condition can be fixed through an equation for the function *p*_*cure*_(*t*) (see Eq. (22)), which is derived using the backward Kolmogorov approach. In Section 5.1, the large-time limit of *p*_*cure*_(*t*) is obtained in closed form in terms of the initial condition. The single-absorbing-point boundary condition above can be defined more rigorously by setting 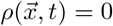 on the line *x* + *y* = *ε*, where *ε «* 1*/N* is sufficiently small (i.e., this line can be defined as an absorbing boundary through which the probability flux gets into a small region near the origin), and by *defining*

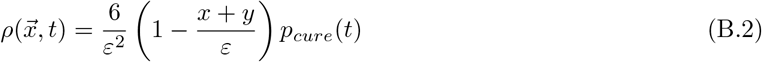

for any (*x, y*) within the small region *{*(*x, y*)*|x ≥* 0; *y ≥* 0; *x* + *y ≤ ε}*. When the probability density (B.2) is integrated over this area, we get exactly *p*_*cure*_(*t*), i.e., the small region near the origin approximately represents the cure state. In the weak limit (in distributional sense) *ε →* 0, the probability density at the origin becomes a Dirac delta distribution.

The boundary condition at the recurrence line *x* + *y* = 1 is also absorbing, i.e., 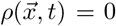. At either *x* = 0 or *y* = 0, the boundary condition for the regular part of 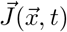 (i.e., the current density defined by Eq. (15), corresponding to the regular part of 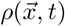) is given by 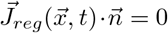, where 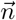 is the outward normal, i.e., there cannot be any flux crossing the boundaries *x* = 0 or *y* = 0, except at the cure state (*x, y*) = (0, 0), at which the regular part of 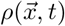 vanishes, whereas the delta peak works as a single absorbing point. The absorbing boundary conditions should be interpreted as follows: while the single absorbing point at the origin pins any probability that it absorbs to the cure state (thus the delta peak), the probability flux through the recurrence boundary *x* + *y* = 1 exits to the outer region *{*(*x, y*)*|x ≥* 0; *y ≥* 0; *x* + *y ≥* 1*}* and never returns.

This proper definition of the boundary condition near the origin ensures that the Fokker-Planck equation (14) gives the correct probability conservation equation in its integral form. Indeed, using the boundary-condition scheme described above and integrating Eq. (14) over the area Ω_*ε*_ = *{*(*x, y*)*|x ≥* 0; *y ≥* 0; *ε ≤ x* + *y ≤* 1*}*, we get

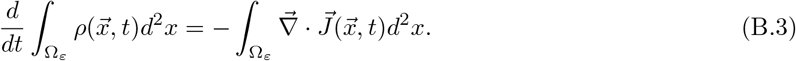

Using the divergence theorem and the reflecting boundary condition 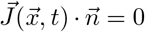 at the boundaries *x* = 0, *ε* ≤ *y* ≤ 1 and *y* = 0, *ε ≤ x ≤* 1, we get

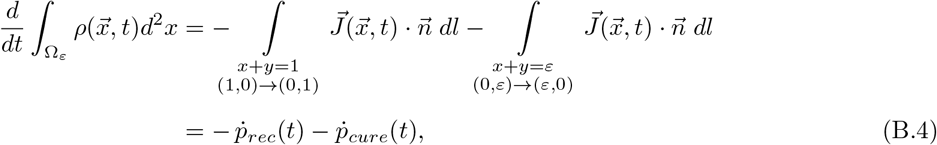

where 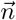 is the outward normal and *p*_*rec*_(*t*), *p*_*cure*_(*t*) are the probabilities of recurrence and cure at any time *≤ t*, respectively.

### Appendix B.2. Backward Kolmogorov equation

The boundary conditions for the backward equation (17) can be derived from those of the forward equation (14) as follows (see e.g. [44]). Let *f* and *g* be arbitrary square-integrable functions defined on the domain Ω_*ε*_ = *{*(*x, y*)*|x ≥* 0; *y ≥* 0; *ε ≤ x* + *y ≤* 1*}*, satisfying the forward and the backward equations/boundary conditions, respectively. Let us consider the *L*^2^ inner product

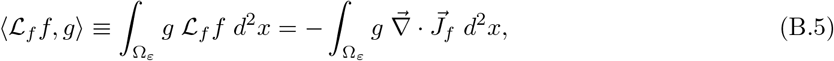

where 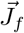 is the current density as defined in Eq. (15) for the density function *f*. Integrating the right-hand side of Eq. (B.5) by parts, it can be shown that

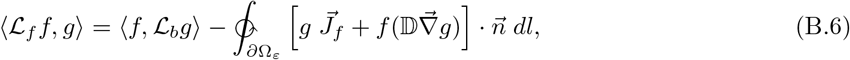

where 𝔻 is the diffusion tensor defined in Eq. (18).

Since the backward operator is the adjoint of the forward operator, i.e., 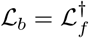, we must have 〈𝓛_*f*_ f, g〉 = 〈f, 𝓛_*b*_g〉 for any square-integrable functions *f, g* defined on the domain Ω_*ε*_ that satisfy the forward and backward equation/boundary conditions, respectively. Therefore, it follows that the boundary term on the right-hand side of Eq. (B.6) must vanish for any such *f, g*. This means that given boundary conditions on any function *f* satisfying the forward equation, the boundary conditions for any function *g* satisfying the backward equation have to be chosen in such a way that the integrand on the second term of Eq. (B.6) must vanish. It then follows that *g* = 0 for absorbing boundaries and 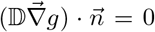 for reflecting boundaries. Hence, on both absorbing boundaries *x*_0_ + *y*_0_ = *ε* and *x*_0_ + *y*_0_ = 1, the boundary condition is 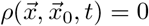. Using Eqs. (17) and (18), we can also show that

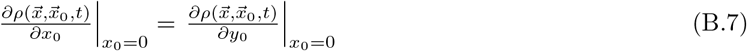

at the reflecting boundary *x*_0_ = 0, *ε ≤ y*_0_ *≤* 1 and

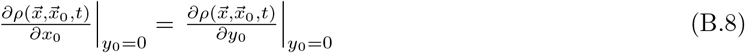

at the reflecting boundary *y*_0_ = 0, *ε ≤ x*_0_ *≤* 1.

## Appendix C. Probability flux through the absorbing boundaries

The probability flux through the recurrence boundary can be obtained by integrating the normal component of the current density (15) over the line *x* + *y* = 1,

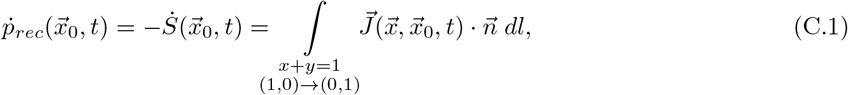

where 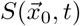 is the recurrence-free survival function (see Eqs. (16) and (B.4)) and 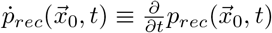. For *µ*_*D*_ = 0, using the boundary condition that the probability density has to vanish at the recurrence line *x* + *y* = 1, we get

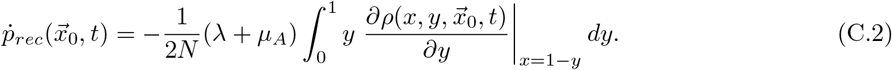

Using the scheme described in Appendix B.1 for the boundary condition near the origin (see discussion below Eq. (B.1) and also Eq. (B.4)), for *µ*_*D*_ = 0 we find that the probability flux into the cure state (0, 0) is given by

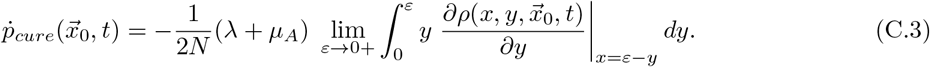

The partial differential equations (21) and (22) for the time-dependent probabilities of recurrence and cure before time *t* can immediately be derived, respectively, by doing the operations on the right-hand sides of Eqs. (C.2) and (C.3) on both sides of Eq. (17). These operations commute with the backward operator 𝓛_*b*_ defined in Eq. (17), because the latter only acts on the initial-condition variables (*x*_0_*, y*_0_). Eqs. (21) and (22) then follow after integration in time from 0 to *t*.

## Appendix D. Solution of the mean recurrence time equation outside the boundary layers (leading order)

At leading (zeroth) order in 1*/N* Eq. (29) becomes the first-order PDE (36), which is valid outside boundary layers that exist near the reflecting boundaries *z*_0_ = 0 and *z*_0_ = *w*_0_. Eq. (36) can be solved by the method of characteristics, as follows.

Along the characteristic curves (which are parametrized by a parameter *r*), the PDE (36) becomes the set of ODEs

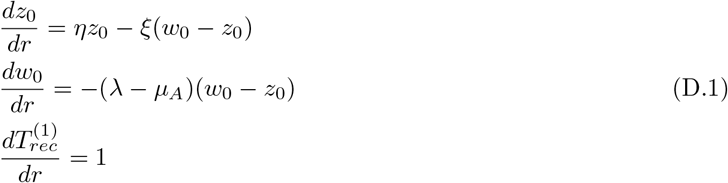

This last equation immediately gives 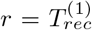 (here we can choose the constant of integration to be zero, since it can be absorbed into the constants *s*_1_ and *s*_2_ in Eqs. (D.2) below).

Solving the first two ODEs yields

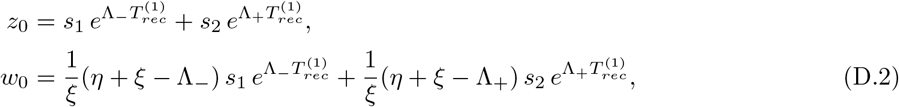

where *s*_1_ and *s*_2_ are constants of integration and

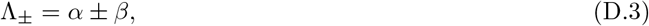

with *α* and *β* given by Eqs. (39) and (40).

By imposing the absorbing boundary condition 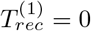 at *w*_0_

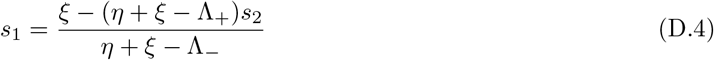

Using this relation in Eqs. (D.2), it follows that

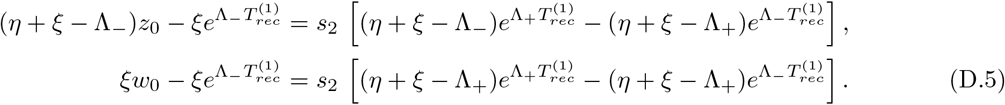

Eliminating the constant of integration *s*_2_ from Eqs. (D.5) and simplifying the resulting equation, we finally get the solution given by Eqs. (37) and (38).

## References

[1] American Cancer Society. Cancer Facts & Figures 2018. Atlanta, American Cancer Society (2018).

[2] P. M. Altrock, L. L. Liu; F. Michor. The mathematics of cancer: integrating quantitative models. Nat. Rev. Cancer 15, 730–745 (2015).

[3] A. Swierniak; A. Polanski; M. Kimmel. Optimal control problems arising in cell-cycle-specific cancer chemotherapy. Cell Prolif. 29, 117–139 (1996).

[4] A. Wu; D. Liao; V. Kirilin; et al. Cancer dormancy and criticality from a game theory perspective. Cancer Converg. 2, 1 (2018).

[5] N. Komarova. Stochastic modeling of drug resistance in cancer. J. Theor. Biol. 239(3), 351–366 (2006).

[6] J. Foo; F. Michor. Evolution of resistance to anti-cancer therapy during general dosing schedules. J. Theor. Biol. 263(2), 179–188 (2010).

[7] T. Antal; P. L. Krapivsky. Exact solution of a two-type branching process: models of tumor progression. J. Stat. Mech. Theory Exp. P08018 (2011).

[8] N. L. Komarova; D. Wodarz. Stochastic modeling of cellular colonies with quiescence: An application to drug resistance in cancer. Theor. Popul. Biol. 72(4), 523–538 (2007).

[9] B. G. Birkhead; E. M. Rankin; S. Gallivan; L. Dones; R. D. Rubens. A mathematical model of the development of drug resistance to cancer chemotherapy. Eur. J. Cancer Clin. Oncol. 23, 1421–1427 (1987).

[10] J. C. Panetta; J. Adam. A mathematical model of cycle-specific chemotherapy. Math. Comput. Modelling 22(2), 67–82 (1995).

[11] J. C. Panetta. A mathematical model of breast and ovarian cancer treated with paclitaxel. Math. Biosci. 146(2), 89–113 (1997).

[12] J. West; P. K. Newton. Chemotherapeutic Dose Scheduling Based on Tumor Growth Rates Provides a Case for Low-Dose Metronomic High-Entropy Therapies. Cancer Res. 77(23), 6717–6728 (2017).

[13] V. T. DeVita; P. S. Schein. The use of drugs in combination for the treatment of cancer: rationale and results. N. Engl. J. Med., 288(19):998–1006 (1973).

[14] S. Benzekry; C. Lamont; A. Beheshti; A. Tracz; J. M. L. Ebos; L. Hlatky; et al. Classical Mathematical Models for Description and Prediction of Experimental Tumor Growth. PLoS Comput. Biol. 10, e1003800 (2014).

[15] L. Norton; R. Simon; H. D. Brereton; A. E. Bogden. Predicting the course of Gompertzian growth. Nature 264, 542–545 (1976).

[16] S. E. Clare; F. Nakhlis; J. C. Panetta. Molecular biology of breast cancer metastasis: The use of mathematical models to determine relapse and to predict response to chemotherapy in breast cancer. Breast Cancer Res. 2(6), 430–435 (2000).

[17] M. W. Retsky; D. E. Swartzendruber; R. H. Wardwell; P. D. Bame. Is Gompertzian or exponential kinetics a valid description of individual human cancer growth? Med. Hypotheses 33(2), 95–106 (1990).

[18] M. W. Retsky; D. E. Swartzendruber; P. D. Bame; R. H. Wardwell. Computer Model Challenges Breast Cancer Treatment Strategy. Cancer Investig. 12(6), 559–567 (1994).

[19] J. F. Speer; V. E. Petrosky; M. W. Retsky; R. H. Wardwell. A Stochastic Numerical Model of Breast Cancer Growth That Simulates Clinical Data. Cancer Res. 44, 4124–4130 (1984).

[20] M. W. Retsky; R. Demicheli; D. E. Swartzendruber; P. D. Bame; R. H. Wardwell; G. Bonadonna; J. F. Speer; P. Valagussa. Computer simulation of a breast cancer metastasis model. Breast Cancer Res. Treat. 45, 193–202 (1997).

[21] R. Demicheli; R. Miceli; C. Brambilla; L. Ferrari; A. Moliterni; M. Zambetti; P. Valagussa; G. Bonadonna. Comparative analysis of breast cancer recurrence risk for patients receiving or not receiving adjuvant cyclophosphamide, methotrexate, fluorouracil (CMF). Data supporting the occurrence of ‘cures’. Breast Cancer Res. Treat. 53, 209–215 (1999).

[22] P. Alexander. 2nd Gordon Hamilton Fairley lecture. Need for new approaches to the treatment of patients in clinical remission, with special reference to acute myeloid leukaemia. Br. J. Cancer 46, 151–159 (1982).

[23] Early Breast Cancer Trialists’ Collaborative Group (EBCTCG). Adjuvant chemotherapy in oestrogen-receptor-poor breast cancer: patient-level meta-analysis of randomised trials. Lancet, vol. 371, issue 9606, pp. 29–40 (2008).

[24] D. F. Hayes; A. D. Thor; L. G. Dressler; et al. HER2 and Response to Paclitaxel in Node-Positive Breast Cancer. N. Engl. J. Med., 357:1496–1506 (2007).

[25] N. W. Wilkinson; G. Yothers; S. Lopa; J. P. Costantino; N. J. Petrelli; N. Wolmark. Long-Term Survival Results of Surgery Alone Versus Surgery Plus 5-Fluorouracil and Leucovorin for Stage II and Stage III Colon Cancer: Pooled Analysis of NSABP C-01 Through C-05. A Baseline from Which to Compare Modern Adjuvant Trials. Ann. Surg. Oncol., Vol. 17, 959–966 (2010).

[26] D. A. Berry; C. Cirrincione; I. C. Henderson; et al. Estrogen-receptor status and outcomes of modern chemotherapy for patients with node-positive breast cancer. JAMA 295: 1658–1667 (2006).

[27] K. Dunleavy; S. Pittaluga; M. Shovlin; et al. Low-Intensity Therapy in Adults with Burkitt’s Lymphoma. N. Engl. J. Med., 369:1915–1925 (2013).

[28] W. J. Gradishar; R. Hellmund. A Rationale for the Reinitiation of Adjuvant Tamoxifen Therapy in Women Receiving Fewer than 5 Years of Therapy. Clin. Breast Cancer, Vol. 2, Issue 4, pp. 282–286 (2002).

[29] N. Gautam. Analysis of Queues: Methods and Applications. CRC Press (2012).

[30] G. Latouche; V. Ramaswami. Introduction to Matrix Analytic Methods in Stochastic Modeling. ASA-SIAM Series on Statistics and Applied Probability. SIAM, Philadelphia, PA (1999).

[31] R. A. Usmani. Inversion of Jacobi’s tridiagonal matrix. Computers Math. Applic., Vol. 27, No. 8, pp. 59–66 (1994).

[32] Y. Huang; W. F. McColl. Analytical inversion of general tridiagonal matrices. J. Phys. A: Math. Gen. 30, 7919 (1997).

[33] J. P. Kharoufeh. Level-dependent quasi-birth-and-death processes. Wiley Encyclopedia of Operations Research and Management Science, John Wiley & Sons, Inc., Hoboken, NJ (2011).

[34] V. Ramaswami; P. G. Taylor. Some properties of the rate operators in level dependent quasi-birth-and-death processes with a countable number of phases. Stochastic Models 12, 143–164 (1996).

[35] L. Bright; P. G. Taylor. Calculating the equilibrium distribution in level dependent quasi-birth-and-death processes. Stochastic Models 11(3), 497–525 (1995).

[36] L. S. T. Ho; J. Xu; F. W. Crawford; V. N. Minin; M. A. Suchard. Birth/birth-death processes and their computable transition probabilities with biological applications. J. Math. Biol. 76: 911–944 (2018).

[37] J. Abate; W. Whitt. Transient behavior of the M/M/1 queue via Laplace transforms. Adv. Appl. Prob. 20, 145–178 (1988).

[38] J. Abate; W. Whitt. Transient behavior of the M/M/1 queue: Starting at the origin. Queueing Syst. 2, 41 (1987).

[39] J. Abate; W. Whitt. Approximations for the M/M/1 busy-period distribution. In: Queueing Theory and its Applications, Liber Amicorum Professor J. W. Cohen, O. J. Boxma and R. Syski (eds.) North-Holland, Amsterdam, 149–191 (1988).

[40] P. Leguesdron; J. Pellaumail; G. Rubino; B. Sericola. Transient analysis of the M/M/1 queue. Adv. Appl. Prob. 25, 702–713 (1993).

[41] O. P. Sharma; B. D. Bunday. A Simple Formula for the Transient State Probabilities of an M/M/1/∞ Queue. Optimization 40, 79–84 (1997).

[42] O. P. Sharma; N. S. K. Nair. Transient analysis of finite state birth and death process with absorbing boundary states. Stoch. Anal. Appl. 14:5, 565–589 (1996).

[43] H. Risken. The Fokker–Planck equation: methods of solution and applications. Berlin, Springer (1984).

[44] C. W. Gardiner. Handbook of Stochastic Methods for Physics, Chemistry and the Natural Sciences. Berlin, Springer (1985).

[45] S. Iyer-Biswas; A. Zilman First-passage processes in cellular biology. Adv. Chem. Phys. 160, 261 (2016).

[46] G. H. Weiss. First passage time problems in chemical physics. Adv. Chem. Phys. 13, 1–18 (1967).

[47] T. Chou; M. R. D’Orsogna. First Passage Problems in Biology. In: R. Metzler, G. Oshanin and S. Redner (eds). First-Passage Phenomena and Their Applications. World Scientific, pp. 306–345 (2014).

[48] S. Redner. A Guide to First-Passage Processes. Cambridge, UK, Cambridge Univ. Press (2001).

[49] M. H. Holmes. Introduction to Perturbation Methods. Springer-Verlag, New York (1995).

[50] Z. Liu; H. Lou; K. Xie; H. Wang; N. Chen; O. M. Aparicio; M. Q. Zhang; R. Jiang; T. Chen. Reconstructing cell cycle pseudo time-series via single-cell transcriptome data. Nat. Commun. 8, 22 (2017).

[51] Z. Ji; H. Ji. TSCAN: Pseudo-time reconstruction and evaluation in single-cell RNA-seq analysis. Nucleic Acids Res. 44(13), e117 (2016).

[52] M. S. Kowalczyk; I. Tirosh; D. Heckl; et al. Single cell RNA-seq reveals changes in cell cycle and differentiation programs upon aging of hematopoietic stem cells. Genome Res. 25, 1860–1872 (2015).

[53] C. R. Doering; K. V. Sargsyan; L. M. Sander. Extinction Times for Birth-Death Processes: Exact Results, Continuum Asymptotics, and the Failure of the Fokker-Planck Approximation. Multiscale Model. Simul. 3(2), 283–299 (2005).

[54] The Cancer Genome Atlas: https://cancergenome.nih.gov, https://portal.gdc.cancer.gov

[55] F. W. Crawford, L. S. T. Ho, M. A. Suchard. Computational methods for birth-death processes. WIREs Comput Stat., e1423 (2018).

[56] E. L. Kaplan; P. Meier. Nonparametric Estimation from Incomplete Observations. J. Am. Stat. Assoc. 53(282), 457–481 (1958).

[57] L. Y. Han; V. Karavasilis; T. v. Hagen; et al. Doubling time of serum CA125 is an independent prognostic factor for survival in patients with ovarian cancer relapsing after first-line chemotherapy. Eur. J. Cancer 46(8), 1359–1364 (2010).

